# Allosteric Regulation of the EphA2 Receptor Intracellular Region by Serine/Threonine Kinases

**DOI:** 10.1101/2021.01.28.428700

**Authors:** Bernhard C. Lechtenberg, Marina P. Gehring, Taylor P. Light, Mike W. Matsumoto, Kalina Hristova, Elena B. Pasquale

## Abstract

Eph receptor tyrosine kinases play a key role in cell-cell communication. However, lack of structural information on the entire multi-domain intracellular region of any Eph receptor has hindered detailed understanding of their signaling mechanisms. Here, we use an integrative structural biology approach combining X-ray crystallography, small-angle X-ray scattering and hydrogen-deuterium exchange mass spectrometry, to gain the first insights into the structure and dynamics of the entire EphA2 intracellular region. EphA2 promotes cancer malignancy through a poorly understood non-canonical form of signaling that depends on serine/threonine phosphorylation of the linker connecting the EphA2 kinase and SAM domains. We uncovered two distinct molecular mechanisms that may function in concert to mediate the effects of linker phosphorylation through an orchestrated allosteric regulatory network. The first involves a shift in the equilibrium between a “closed” configuration of the EphA2 intracellular region and an “open” more extended configuration induced by the accumulation of phosphorylation sites in the linker. This implies that cooperation of multiple serine/threonine kinase signaling networks is necessary to promote robust EphA2 non-canonical signaling. The second involves allosteric rearrangements in the kinase domain and juxtamembrane segment induced by phosphorylation of some linker residues, suggesting a link between EphA2 non-canonical signaling and canonical signaling through tyrosine phosphorylation. Given the key role of EphA2 in cancer malignancy, this new knowledge can inform therapeutic strategies.

## INTRODUCTION

The Eph receptors are the largest of the receptor tyrosine kinase families, and together with their cell surface anchored ephrin ligands represent an important cell-cell communication system that regulates a multitude of physiological and pathological processes^1–4^. The fourteen Eph receptors have a conserved domain structure, with an extracellular portion including the N-terminal ligand-binding domain and several other domains^5,6^. Following the transmembrane helix, the intracellular portion of the Eph receptors includes a juxtamembrane segment of ~50 amino acids, the tyrosine kinase domain, a ~20 amino acid-long linker, a sterile alpha motif (SAM) domain, and a short C-terminal tail containing a PDZ domain-binding motif (Fig. 1A). The juxtamembrane segment can control kinase activity through a mechanism involving tyrosine phosphorylation^7–9^. Effects of the SAM domain on receptor kinase activity have also been reported, which may differ depending on the Eph receptor^10–13^. How the SAM domain affects Eph receptor kinase activity and the potential interplay between the juxtamembrane segment and SAM domain are poorly understood. While structural information on the entire multidomain extracellular portion of Eph receptors has provided important insights^5,14^, lack of structural information on the configuration of the entire intracellular region of any Eph receptor has hindered detailed understanding of their signaling mechanisms.

**Figure 1.**
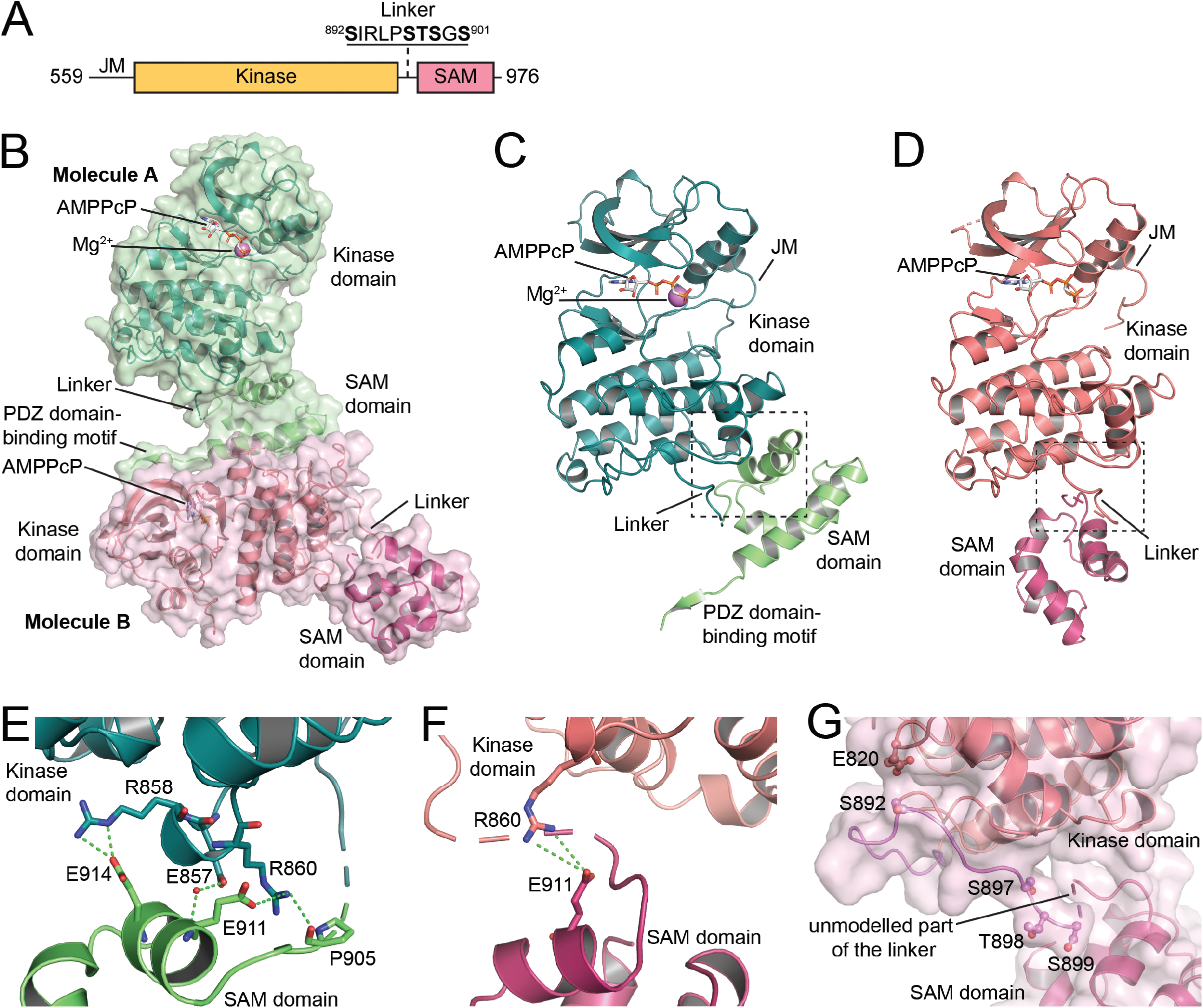
Crystal structure of the EphA2 wild-type intracellular region. (**A**) Schematic illustrating the domains of the EphA2 intracellular region. (**B**) Overview of the two EphA2 molecules in the crystallographic asymmetric unit. Molecule A is shown in green, with the kinase domain in darker green and the SAM domain and C-terminal tail in lighter green. The kinase and SAM domains of molecule B are shown in two different shades of red. The ATP-analogue AMPPCP, present in the active sites of both molecules A and B, is shown as grey sticks and a Mg^2 +^-ion in the active site of molecule A is shown as a purple sphere. The kinase domains in both molecules adopt the active DFG-in conformation. (**C**) Rotated view of molecule A, highlighting the compact arrangement of the kinase and SAM domains. The dashed box indicates the region highlighted in panel E. (**D**) Rotated view of molecule B, with the kinase domain in the same orientation as for molecule A in panel C. The kinase and SAM domains are in a more elongated arrangement. The dashed box indicates the region highlighted in panel F. (**E**) Detail of the intramolecular interactions between the kinase and SAM domains of molecule A. Interacting residues are shown as sticks with interactions indicated as dashed lines. A water molecule forming hydrogen bonds with the sidechain of E857 and the backbone of E911 is shown as a small red sphere. (**F**) Intramolecular interactions between the kinase and SAM domains of molecule B. (**G**) Detail of the kinase-SAM linker of molecule B, showing that the S892, S897, T898, and S899 phosphorylation sites are solvent exposed and accessible. The fifth phosphorylation site (S901) is in the undefined region of the linker, which is indicated by a dashed connector. E820 is in close proximity (3–5 Å) to S892 and might be an allosteric sensor of linker phosphorylation.

The EphA2 receptor has been implicated in many physiological and pathological processes, including epithelial homeostasis, immune system function, angiogenesis, inflammation, atherosclerosis, parasitic infections, and cancer malignancy^2–4,15,16^. EphA2 can regulate such a wide variety of diverse processes at least in part because it can signal through different mechanisms. EphA2 canonical signaling is typical for a receptor tyrosine kinase; it is induced by ephrin ligand binding and is mediated by kinase activity. EphA2 canonical signaling in tumor endothelial cells promotes tumor angiogenesis^2,17–19^. In cancer cells, however, EphA2 tyrosine phosphorylation is often low, consistent with the unusual effects of EphA2 canonical signaling, which inhibits major oncogenic networks such as RAS-ERK and AKT-mTORC1^2,20,21^. EphA2 can also signal through a completely different, more recently discovered non-canonical signaling mechanism that depends on phosphorylation on serine 897 (S897). EphA2 non-canonical signaling is often elevated in cancer cells and promotes their oncogenic properties, including epithelial-mesenchymal transition, invasiveness and metastasis, mechanotransduction, stem cell-like properties, and drug resistance^2,3,22–25^. Despite its profound significance in cancer, the mechanism of non-canonical signaling is not well understood^22,26^.

S897 is located in the linker connecting the kinase and SAM domains, and it is not known how its phosphorylation can regulate EphA2 signaling properties. For example, the phosphorylated S897 motif does not match consensus binding motifs for protein domains recognizing phosphoserine sites^27^. However, the kinase-SAM linker contains a cluster of 5 serine/threonine residues that in addition to S897 also include S892, T898, S899 and S901. These residues can also be phosphorylated^28,29^(phosphosite.org), but it is not known whether their phosphorylation has functional importance and plays a role in EphA2 non-canonical signaling.

To understand the interplay of the different domains and linkers in the EphA2 intracellular region and mechanistically elucidate how phosphorylation of the kinase-SAM linker affects EphA2 function, we sought to obtain structural information on the configuration of the entire EphA2 intracellular portion. Successful expression of most of the EphA2 intracellular region in soluble form enabled us to solve its crystal structure and investigate its dynamic behavior in solution using small-angle X-ray scattering (SAXS) and hydrogen-deuterium exchange in combination with mass spectrometry (HDX-MS). The data obtained by applying this integrative structural biology approach reveal an allosteric network connecting the kinase-SAM linker with the kinase domain, activation loop and juxtamembrane segment. Furthermore, they support a model in which multiple negative charges introduced by linker phosphorylation cooperate to promote an “open” conformation of the EphA2 intracellular region that mediates non-canonical signaling. We indeed found that multiple EphA2 kinase-SAM linker residues can be simultaneously phosphorylated by different kinases, consistent with the notion that not only the single negative charge introduced by S897 phosphorylation but multiple phosphorylated residues in the linker are important for EphA2 non-canonical signaling.

## RESULTS

### S897 is part of a cluster of phosphorylation sites in the EphA2 kinase-SAM linker

Mass spectrometry data from the PhosphoSitePlus database show that all five serine/threonine residues in the EphA2 kinase-SAM linker are frequently phosphorylated in cell lines and tissues (Fig. S1A). By screening a number of cancer cell lines in which EphA2 is known to be phosphorylated on S897^28,30^, we confirmed that in these cell lines EphA2 is also phosphorylated on two other residues in the cluster, S892 and S901, as determined by immunoblotting with phosphospecific antibodies we generated^28,29^(Fig. S1B,C). We observed a strong correlation between the extent of EphA2 phosphorylation on S897 and S901 (Fig. S1D), consistent with our previous findings that EphA2 phosphorylation on S901 depends on the negative charge introduced by phosphorylation on S897^29^. Furthermore, our analysis of published mass spectrometry data profiling 27 non-small cell lung cancer and 4 breast cancer cell lines^30^ shows a very strong correlation of EphA2 phosphorylation on S897 with phosphorylation on T898, S899 and S901 and a weaker correlation with S892 phosphorylation (Fig. S1E). Consistent with these findings, mass spectrometry data show that more than 60% of the EphA2 kinase-SAM linker peptides analyzed are phosphorylated on multiple residues simultaneously (Fig. S1F)^28,31—36^. Up to three phosphorylation sites have been detected in the same EphA2 linker peptide, which could be an underestimate since some dephosphorylation likely occurs during sample preparation. Thus, EphA2 kinase-SAM linker phosphorylation is highly cooperative, and S897 is typically phosphorylated together with other linker residues.

### Crystal structure of the EphA2 intracellular region including the kinase and SAM domains

To obtain insights into the configuration of the EphA2 intracellular region and its role in EphA2 signaling, we solved the crystal structure of the entire unphosphorylated EphA2 intracellular region except for the first 30 residues (residues D590-I976), bound to the ATP analog β,γ-methyleneadenosine 5′-triphosphate (AMPPcP), at a resolution of 1.75 Å (Fig. 1, Table S1). To our knowledge, this represents the first structure of the almost entire intracellular region of any Eph receptor and provides novel insights into the kinase-SAM domain arrangement.

The crystallographic asymmetric unit contains two EphA2 molecules (molecules A and B; Fig. 1B-D), even though the EphA2 intracellular portion is monomeric in solution, as shown by analytical ultracentrifugation (Fig. S2). The regions that are well defined in both EphA2 molecules include the portion of the juxtamembrane segment present in our construct, most of the kinase domain, and the whole SAM domain (Fig. 1B). However, parts of the activation loop (residues L760-I779), residues S636-G637 in the short β1-2 loop of the N-lobe of EphA2 molecule B, and parts of the kinase-SAM linker (residues S899-V904 in molecule A and G900-V904 in molecule B) are not defined due to lack of electron density (Fig. 1C,D). Furthermore, the C-terminal tail (K966-I976) including the PDZ domain-binding motif is defined only in molecule A (Fig. 1B,C).

Interestingly, residues D590-A600 in the juxtamembrane segment adopt a different conformation compared to previous structures of EphA2 and other Eph receptors containing the juxtamembrane segment and kinase domain. In our structure, the αA′ helix is unwound and Y594 (the second of the two conserved tyrosines in the juxtamembrane segment) occupies a pocket usually occupied by the first conserved tyrosine (Y588 in EphA2, which is not part of our construct) (Fig S3A,B)^9,37^. This observation suggests a dynamic conformation of the juxtamembrane segment even when its tyrosines are unphosphorylated.

The relative orientation of the kinase and SAM domains is different between the two EphA2 molecules. Molecule A has a more compact configuration that is stabilized by a salt bridge and a hydrogen bond formed between the kinase and SAM domains (Fig. 1C,E), whereas in molecule B the two domains are further apart from each other, with only a single weak salt bridge between them (Fig. 1D,F). This suggests a degree of variation in the relative arrangement of the kinase and SAM domains, likely enabled by conformational flexibility in the kinase-SAM linker. The N-terminal part of the linker (up to residue T898, the last residue visible in both EphA2 molecules) as well as the beginning of the SAM domain (starting at F906) adopt the same conformations in both molecules (Fig S3C,D). In contrast, residues S899-P905 (including the S899 and S901 phosphorylation sites) are mostly not defined in both EphA2 molecules, likely due to high conformational flexibility. Thus, conformational differences in these residues appear to be the main reason for the different relative orientations of the kinase and SAM domains in the two EphA2 molecules. The conformational flexibility of the S901 motif also supports the previously proposed idea that the negative charge introduced by S897 phosphorylation, and not a specific conformational change induced by S897 phosphorylation, is critical for subsequent phosphorylation of S901 by CK1 acidophilic kinases^28^.

The N-terminal part of the linker is partly wrapped around the bottom of the kinase domain C-lobe (Fig. 1G) and locked in place by interactions of I893 with a hydrophobic pocket in the kinase domain (Fig S3C). This linker conformation is different from that in previous EphA2 structures that do not contain the SAM domain, where most of the linker (including I893) faces away from the kinase domain (Fig S3C), suggesting that the SAM domain affects linker conformation. On the other hand, the configuration of the N-terminal part of the EphA2 linker and the interaction of I893 with a pocket in the kinase domain resemble the linker configuration in an EphA3 structure lacking the SAM domain, while the central parts of the EphA2 and EphA3 linkers have distinct conformations (Fig S3C)^38^. Importantly, as expected for kinase substrate sites, all serine/threonine residues in the EphA2 linker are fully solvent accessible.

### The arrangement of the EphA2 kinase and SAM domains is regulated by phosphorylation of their linker

To determine whether the negative charges introduced by phosphorylation of the linker affect the kinase-SAM arrangement, we solved the structures of two EphA2 phosphomimetic mutants. In the first mutant, we replaced both S897 and S901 with glutamic acid to mimic the negative charge of the phosphate group. The structure of the EphA2 S897E/S901E double mutant (obtained at a resolution of 2.8 Å) shows a very different kinase-SAM domain arrangement than EphA2 WT (Fig. 2, Table S1). In the single molecule of the crystallographic asymmetric unit, the SAM domain is completely detached from the kinase domain and the C-terminal part of the linker is fully extended (Fig. 2A). We did not observe any substantial differences in the overall fold of the individual domains between the WT and S897E/S901E structures, and all EphA2 molecules in both structures adopt the active DFG-in conformation. Unlike the EphA2 WT structure, the activation loop is completely defined in the S897E/S901E mutant structure, potentially due to stabilizing crystal contacts. Importantly, however, in the S897E/S901E structure the central portion of the kinase-SAM linker adopts a different conformation than that observed in the WT structure and appears to be stabilized by a salt bridge between E897 (replacing S897) and R890 (Fig. 2B). Additionally, we observed a key difference in the αFG loop, including W819 and adjacent residues (Fig. 2B), a region that has previously been identified in EphA3 as part of an allosteric network connecting the juxtamembrane segment, activation loop and SAM linker^39^.

**Figure 2.**
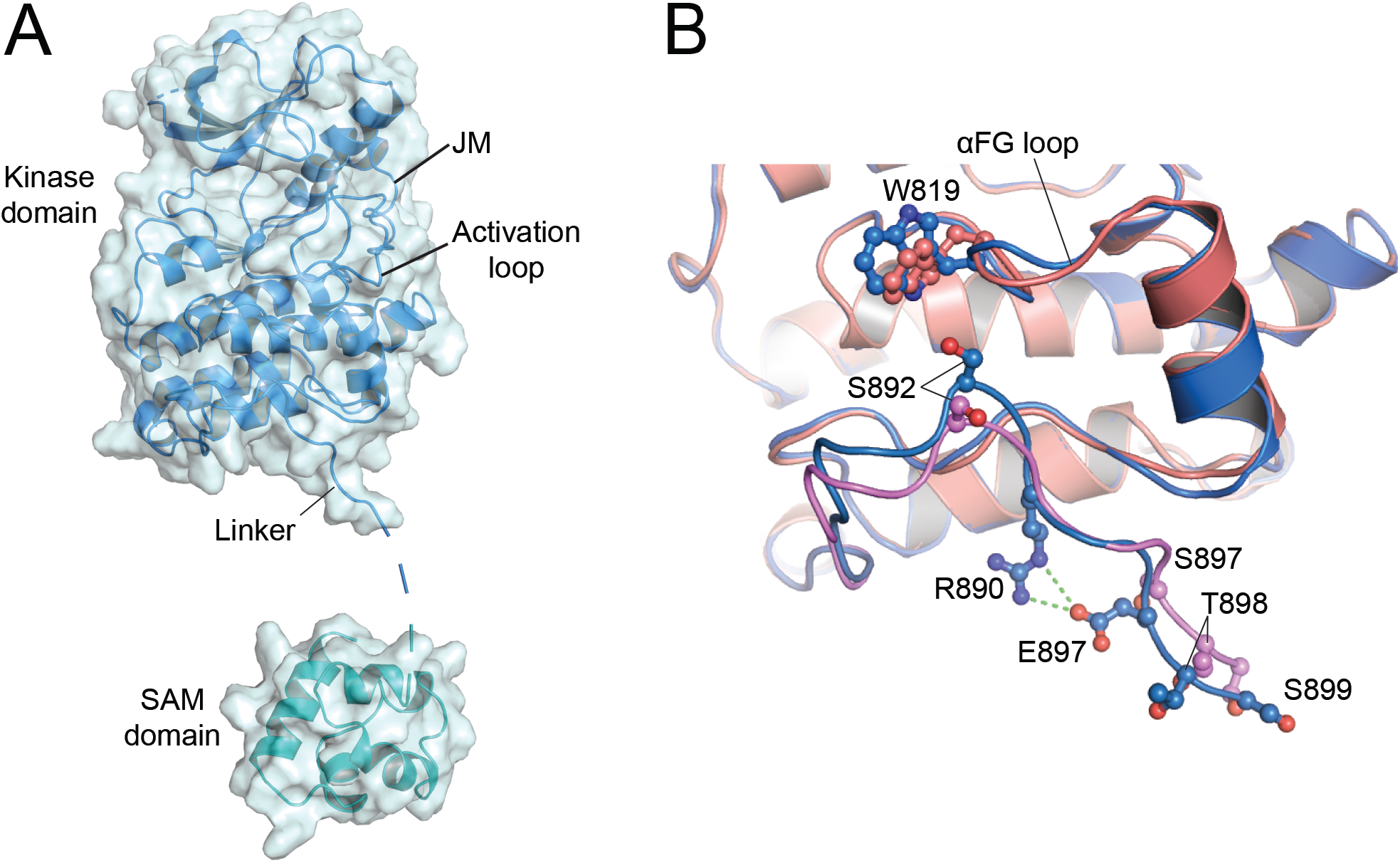
Crystal structure of the intracellular region of the EphA2 S897E/S901E phosphomimetic mutant. (**A**) Overview of the asymmetric unit containing a single EphA2 molecule in a very elongated conformation. The kinase domain is shown in blue and the SAM domain in cyan. The structure includes most of the juxtamembrane segment (JM) and the kinase domain, the full activation loop (including residues L760-I779, which are not defined in the WT structure), and the SAM domain (residuesV909-L965). Parts of the linker region and N-terminus of the SAM domain (residues 900-908, dashed line) and the C-terminal tail (K966-I976) are not visible in the structure due to missing electron density. (**B**) Comparison of the resolved linker regions in the structures of EphA2 WT molecule B (colored as in Fig. 1D) and the S897E/S901E mutant (blue), highlighting key differences in the linker structures. Key residues are shown as sticks and labelled. R890 and the phosphomimetic E897 in the S897E/S901E structure form a salt bridge (green dashes). The SAM domains are omitted for clarity.

In the second EphA2 mutant, we replaced only S901 with glutamic acid. The S901E mutant crystallized under the same conditions as EphA2 WT. In this structure (obtained at a resolution of 2.3 Å), the relative orientations of the kinase and SAM domains are identical to those of EphA2 WT (Fig. S4; Table S1). This is in line with the observation that residue S901 is not defined in the WT crystal structure and suggests that phosphorylation of S901 alone does not significantly affect the conformation of the intracellular portion of EphA2.

Taken together, the data with the two EphA2 mutants suggest that linker phosphorylation regulates the arrangement of the EphA2 kinase and SAM domains and may induce allosteric changes within in the kinase domain. However, since the kinase and SAM domains arrangement in the various crystal structures might be differentially affected by crystal packing, we next used size-exclusion chromatography in line with SAXS (SEC-SAXS) to obtain information on the EphA2 kinase-SAM domain arrangement in solution.

### Solution scattering structure of the EphA2 intracellular region

We first determined the low resolution solution structure of the EphA2 intracellular region (residues D590-I976). Comparison of the SAXS solution scatter data with the scatter data calculated from the crystal structures of EphA2 WT molecules A and B as well as the S897E/S901E double mutant shows that the SAXS data most closely match the conformation observed in EphA2 WT molecule B of the crystallographic asymmetric unit (Fig. 3A). The crystal structure of the EphA2 S897E/S901E mutant shows the most divergent fit to the SAXS solution data (Fig. 3B). The SAXS interatomic distance distribution plot shows a peak with a prominent tail towards longer distances, indicative of an elongated molecule, with a maximum particle dimension (D_max_) of 104 Å (Fig. 3C). Accordingly, ab-initio model building from the SAXS data yields an elongated envelope with a bulky top, likely representing the kinase domain, and a narrower bottom, likely representing the SAM domain (Fig. 3D). Overlay of the SAXS envelope with our EphA2 crystal structures indicates that the SAXS data match best with EphA2 WT molecule B (Fig. 3D), consistent with the calculated scatter data (Fig. 3A-C). However, none of the crystal structures perfectly fits the SAXS envelope, suggesting that the SAXS envelope may represent an ensemble of multiple dynamic conformations of the EphA2 intracellular region in solution. In fact, we did not observe a notable difference between the SAXS curves for EphA2 WT and the EphA2 S897E/S901E double mutant (Fig. 3E), in contrast to the marked differences observed in the crystal structures. We also did not observe notable differences between the SAXS curves for EphA2 WT and the S897E or S901E single mutants (data not shown).

**Figure 3.**
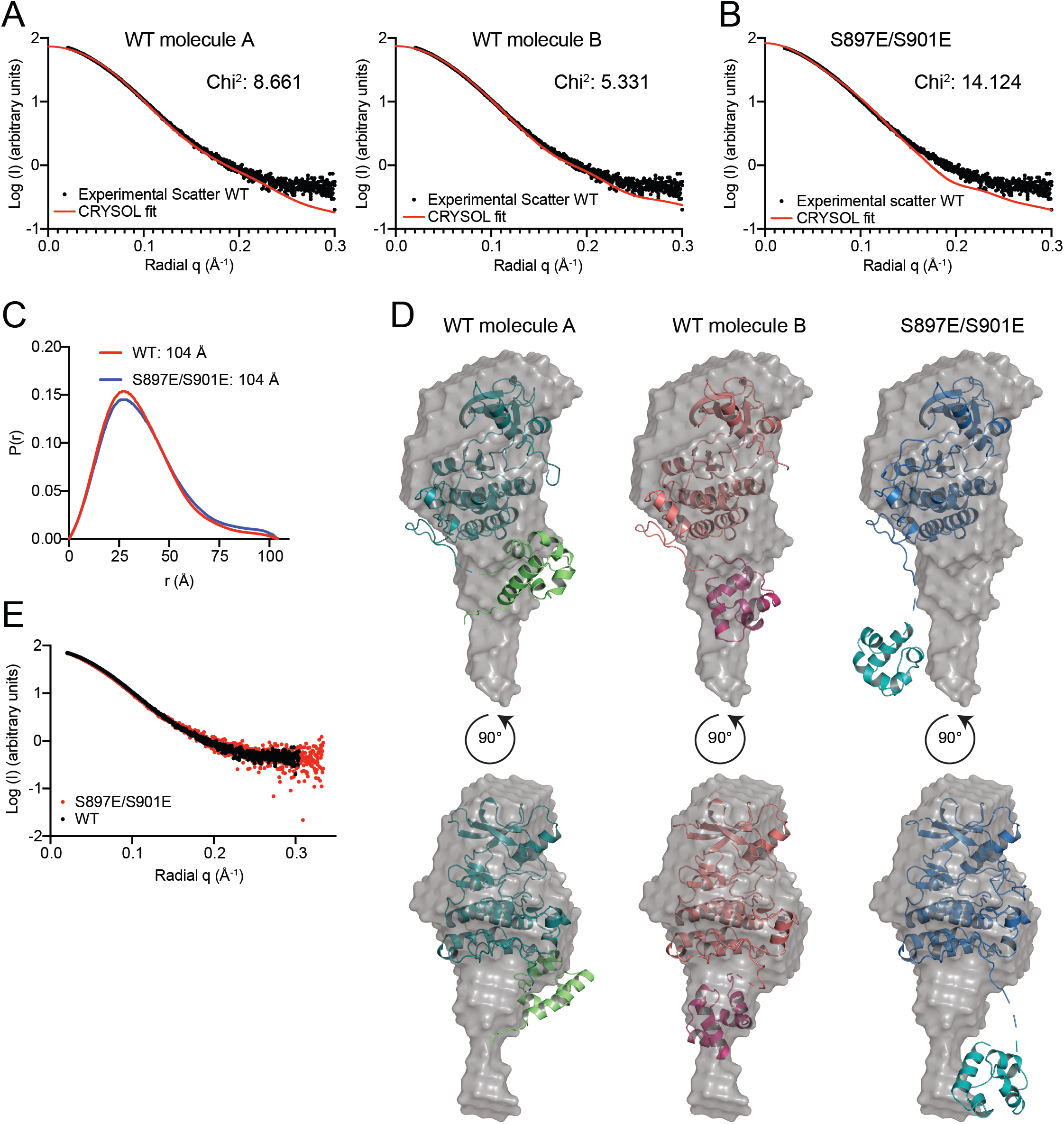
SEC-SAXS analysis of the EphA2 intracellular region. (**A**) Small-angle scattering profile of averaged and background subtracted data from the apex of the in-line size exclusion chromatography peak (black circles) for EphA2 WT. The agreement between the experimental and theoretical scatter calculated from EphA2 molecules A (left) and B (right) using CRYSOL (red line) is reflected in the χ^2^values, indicating that the structure of molecule B more closely resembles the experimental solution scattering data. (**B**) Experimental data for EphA2 WT as in panel A (black circles) shown with the theoretical scatter calculated from the EphA2 S897E/S901E mutant structure using CRYSOL (red line). (**C**) Interatomic distance distribution plot, P(r), calculated from scattering data with GNOM. Maximum particle dimension (D_max_) was estimated as 104 Å for both the EphA2 WT and S897E/S901E mutant intracellular regions. (**D**) Ab initio SAXS model calculation (grey envelope) and fit to molecule A (left) and B (middle) of the EphA2 WT crystal structure and to the EphA2 S897E/S901E mutant structure (right). (**E**) Small-angle scatter profiles, comparing EphA2 WT (black circles, same data as in panels A and B) and the S897E/S901E mutant (red circles).

### Hydrogen-deuterium exchange reveals that the EphA2 kinase domain interacts with the SAM domain

SAXS provides information about the conformation of a protein in solution but, on its own, yields limited detail on protein dynamics. Thus, to further investigate the dynamics of the EphA2 intracellular region, we used HDX-MS to analyze the entire unphosphorylated EphA2 WT intracellular region, except for the first 10 residues (residues S570-I976). This portion of EphA2 comprises 20 additional juxtamembrane residues compared to the portion analyzed by X-ray crystallography and SAXS, and includes the conserved Y588 and Y594, which are known to interact with the kinase domain in unphosphorylated Eph receptors^9^. These HDX-MS experiments identify regions where protein backbone protons in EphA2 rapidly exchange with deuterons from the solvent, indicating that they are more solvent exposed than protons exchanging slowly (Figs. 4, S5, S6). As expected, we found that the core of the kinase domain, including the C-terminal half of the αC helix, the hydrophobic spines and the αF-helix, undergo a small amount of exchange within the 5 min time course of the experiment, indicating that these regions are shielded from the solvent (Fig. 4A). Conversely, the juxtamembrane segment, the Gly-rich loop, the activation loop, and the kinase-SAM linker show rapid exchange even at the earliest time point (0.5 min; Fig. 4A), indicating that these regions are solvent exposed and highly dynamic, in agreement with the crystal structure (Fig. 4B). The solvent exposed, unstructured conformation of the kinase-SAM linker is typical of protein segments containing multiple phosphorylation sites^40^. The high solvent exposure of the juxtamembrane segment suggests that the binding of the conserved juxtamembrane tyrosines to pockets in the kinase domain is not sufficiently stable to detectably alter hydrogen-deuterium exchange, in line with the fact that these tyrosines are phosphorylation sites and thus must be accessible. The observed dynamics of the juxtamembrane segment are also consistent with the discrepancies observed in different crystal structures, which show that Y588 and Y594 can both interact with the same pocket of the EphA2 kinase domain (Fig. S3A,B).

**Figure 4.**
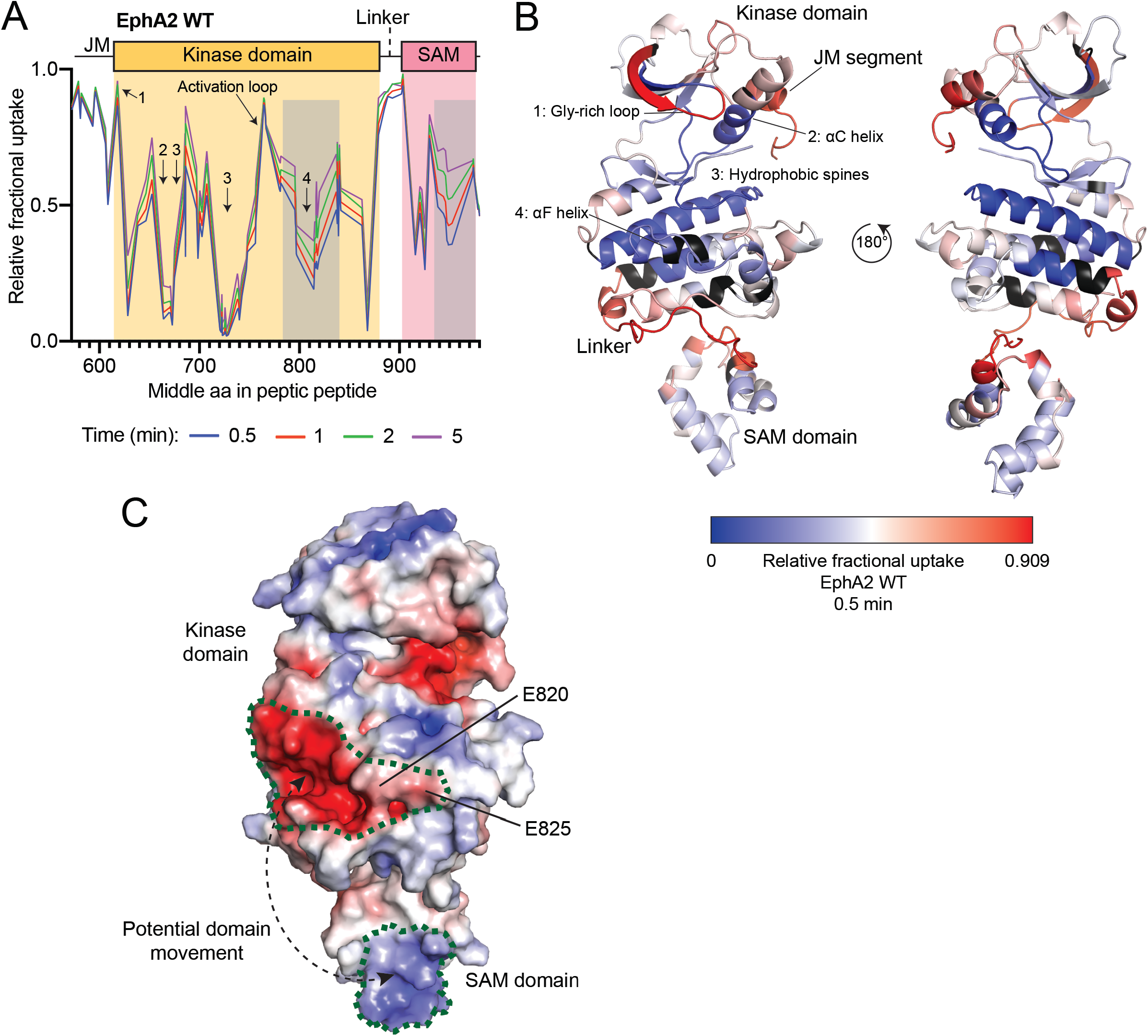
HDX-MS analysis of the EphA2 WT intracellular region. (**A**) Relative fractional deuterium uptake plot for peptic peptides identified from EphA2 WT after 0.5, 1, 2 and 5 min. Key domains of the EphA2 intracellular region are indicated and parts of the kinase domain are indicated with Arabic numerals (see also panel B). The grey shaded areas indicate regions of intermediate exchange, as described in the text. (**B**) Relative fractional deuterium uptake after 0.5 min, mapped onto the EphA2 WT structure (molecule B). Residues without coverage are colored in black. Domains and key regions are indicated. (**C**) Electrostatic surface of the EphA2 WT structure. The positively charged (blue) region in the SAM domain and the negatively charged (red) region in the kinase domain are outlined with a green dotted line. A potential domain movement is indicated by dashed line with arrows. This interaction is possible due to the 20-amino acid flexible linker connecting the SAM and kinase domains. The molecule is shown in the same orientation as that depicted on the left in panel B.

In general, the extent of exchange is similar at the different time points examined. However, in two regions we observed intermediate exchange that gradually increases from 0.5 to 5 min (Fig. 4A,B). One of these regions corresponds to residues P780-M850, comprising the αF–aH helices in the C-lobe of the kinase domain (Fig. 4B). Parts of this region form a prominent negatively charged surface (Fig. 4C). The other region is the C-terminal portion of the SAM domain (residues Y930-N970), which forms a positively charged surface (Fig. 4C).

The correlation of these two exchange processes suggests that the two domains interact with each other through their differently charged surfaces, shielding them from the solvent (Fig. 4B,C). Although we cannot exclude that the two regions are allosterically coupled, this seems unlikely since the C-terminal portion of the linker preceding the SAM domain is highly flexible, as indicated by the rapid hydrogen-deuterium exchange and the lack of electron density in the crystal structures.

### The interaction between the EphA2 kinase and SAM domains is disrupted by negative charges in their linker

We hypothesized that multiple phosphorylation events may lead to effects whose magnitude depends on the number of residues simultaneously phosphorylated. We therefore analyzed the EphA2 5E mutant (S892E/S897E/T898E/S899E/S901E), in which all five phosphosites in the kinase-SAM linker are replaced by glutamic acid, to maximize our ability to detect potential effects of linker phosphorylation. The most drastic effect in this mutant, compared to EphA2 WT, is increased hydrogen-deuterium exchange in the C-lobe of the kinase domain and most of the SAM domain (Figs. 5A,B, S5, S6), suggesting that negative charges in the kinase-SAM linker disrupt the intramolecular interaction between the two domains. Given these results, we compared the hydrogen-deuterium exchange kinetics for EphA2 WT and a set of four different phosphomimetic mutants: 5E, 3E (T898E/S899E/S901E), S892E and S897E (Figs. 5A,C,D,E, S5, S6).

**Figure 5.**
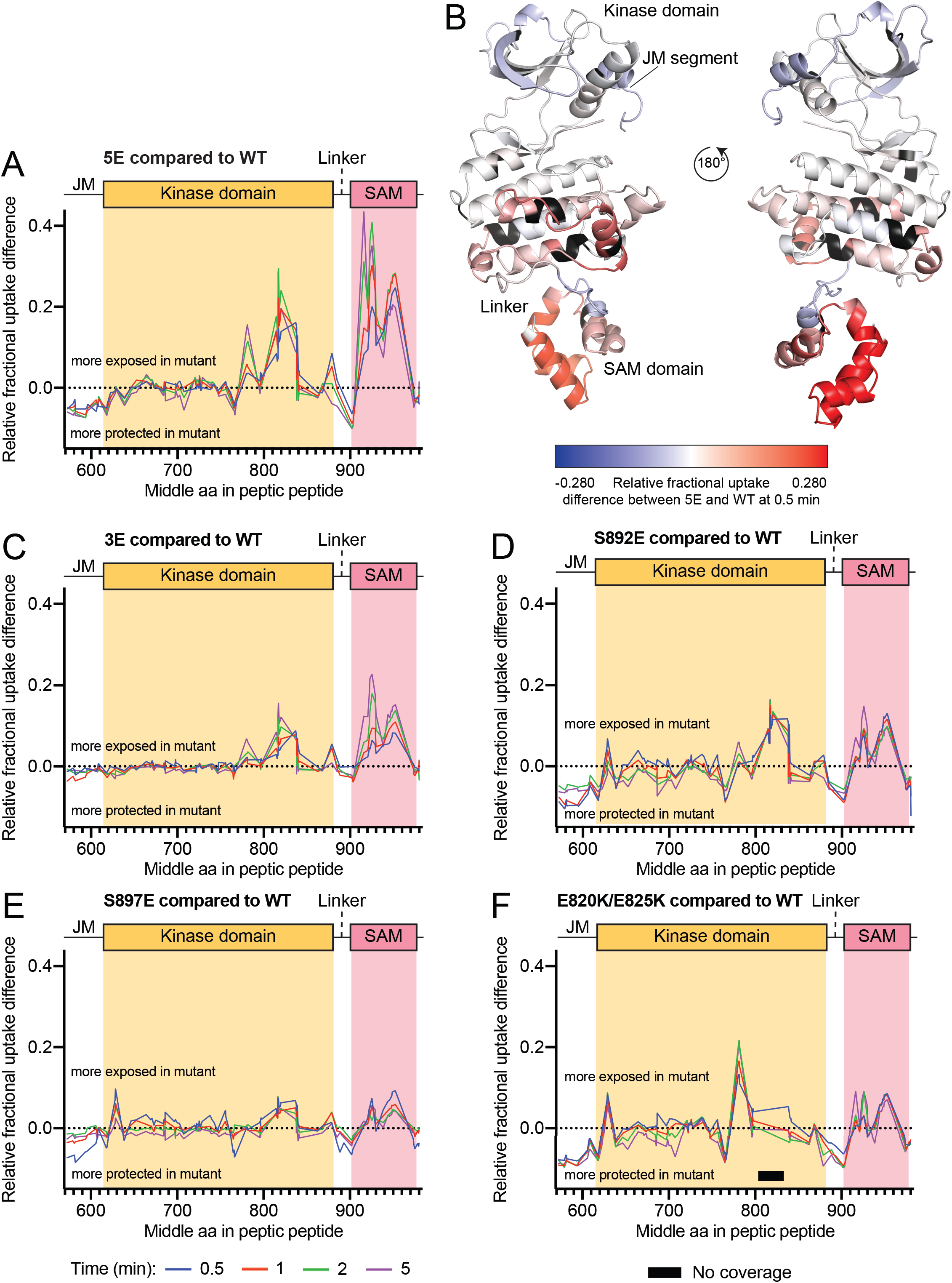
HDX-MS analysis of the EphA2 intracellular region with kinase-SAM linker phosphomimetic mutations. (**A**) Fractional deuterium uptake difference plot comparing the EphA2 5E mutant with EphA2 WT after 0.5, 1, 2 and 5 min of exchange. Positive values indicate more rapid exchange in the 5E mutant compared to WT, whereas negative values indicate slower exchange in the mutant compared to WT. (**B**) Relative fractional deuterium uptake differences between WT and the 5E mutant after 0.5 min mapped onto the EphA2 WT structure (molecule B). Relative fractional differences from −0.183 (less exposed in the 5E mutant) to 0.183 (more exposed in the 5E mutant) are colored in a gradient from blue to red. Residues without coverage are colored in black. (**C-F**) Graphs as in panel A, comparing fractional deuterium uptake differences between the indicated EphA2 mutants and EphA2 WT. The black bar in panel F indicates a region without peptide coverage in the E820K/E825K mutant.

The EphA2 5E mutant shows the most accelerated exchange in the regions of the kinase domain C-lobe and SAM domain that are protected in EphA2 WT (Fig. 5A,B). Surprisingly, the effect of a single negative charge at position 897, the functionally best characterized linker phosphorylation site, does not produce a particularly strong effect on hydrogen-deuterium exchange in these regions (Fig. 5E), suggesting that phosphorylation of S897 alone is not sufficient to regulate EphA2 signaling properties. In fact, a negative charge at position 892 has a stronger effect (Fig. 5D). The three negative residues at positions corresponding to the other three phosphosites in the 3E mutant also accelerate exchange slightly more than the individual S897E mutation (Fig. 5C), suggesting differential effects of the various phosphosites.

We also examined a E820K/E825K mutant, in which we replaced two glutamic acid residues in the negatively charged surface of the kinase domain C-lobe with positively charged lysine residues (Fig. 5F, S5, S6), which we hypothesized would disrupt the observed intramolecular interaction even in the absence of linker phosphorylation. The EphA2 E820K/E825K mutant indeed shows increased exchange in the SAM domain as well as in the region of the kinase domain C-lobe that precedes the introduced mutations (residues P780-V800) (Fig. 5F). Although we did not detect peptides covering residues I805-G833 for this mutant, these observations nevertheless confirm the involvement of the electronegative surface of the EphA2 kinase domain in the interaction with the SAM domain (Fig. 5F).

Interestingly in the EphA2 5E, E820K/E825K and S892E mutants, and to a smaller extent the S897E mutant, we observed a strong correlation in the stabilization of the kinase-SAM linker, parts of the activation loop (residues L760-A770) and the juxtamembrane segment, which undergo lower hydrogen-deuterium exchange in these EphA2 mutants (Fig. 5A,D,E,F). We did not observe this effect in the 3E mutant, in which hydrogen-deuterium exchange in these regions is similar to EphA2 WT (Fig. 5C). This allosteric network may critically depend on E820 in the EphA2 αFG loop, which is located in close proximity to S892 (Fig. 1G).

Together, these data suggest that due to allosteric coupling, phosphorylation of the N-terminal portions of the linker (mostly S892, but also S897) initiates changes in the linker, the activation loop and the juxtamembrane segment, whereas the overall level of phosphorylation in the entire linker controls the interactions between the kinase and SAM domains.

### Phosphorylation of the kinase-SAM linker modestly contributes to EphA2 lateral dimerization in cells

The proposed intramolecular interaction of the SAM domain with the kinase domain in EphA2 molecules positioned side-by-side in the plasma membrane could inhibit their dimerization, possibly by preventing intermolecular interactions between the negatively charged surface of the kinase domain (occupied by the SAM domain) and a positively charged surface on the opposite side of the kinase domain (Fig. S7). We therefore used the fully quantified spectral imaging Forster Resonance Energy Transfer (FSI-FRET) method^41^ to examine how EphA2 mutations mimicking or preventing phosphorylation of kinase-SAM linker residues affect EphA2 lateral interactions in the absence of ligand binding in HEK293T cells. We studied eight different mutants, including 5E, 3E, S897E, S892E, 5A (S892A/S897A/T898A/S899A/S901A), ASAAA (S892A/T898A/S899A/S901A), S897A and S892A, and compared them to EphA2 WT^94^. Homodimerization experiments were performed in cells with EphA2 labeled at the C-terminus with fluorescent proteins (either mTurquoise or EYFP, a FRET pair) attached via a flexible linker (GGS)_5_. FRET efficiencies, which report on the interaction strength between receptor molecules, were measured in the plasma membrane of individual cells, along with the concentration of mTurquoise- and EYFP-labeled EphA2 to obtain dimerization curves (Fig. S8A). Curve fitting yielded the dissociation constants *K_diss_* for the EphA2 mutants, which were compared to K_diss_ for EphA2 WT^94^ (Fig. S8B), showing that the phosphomimetic mutations have negligible effects on EphA2 dimer stability while the non-phosphorylatable alanine mutations appear to have a small destabilizing effect (Fig. S8). The results also suggest effects of the mutations on the geometry of the EphA2 dimers, as inferred from changes in the Intrinsic FRET (Ẽ, Fig. S8B), a structural parameter that depends on the positioning of the fluorescent proteins in the EphA2 dimers. The 3E, S897E, and S897A mutants in particular exhibit higher intrinsic FRET values compared to EphA2 WT (Fig. S8B), which implies that the fluorescent proteins attached to the C-terminus of EphA2 are closer to each other in these dimers.

### Multiple kinase families can phosphorylate the EphA2 kinase-SAM linker

Previous studies have shown that the AGC kinases AKT, RSK and PKA phosphorylate EphA2 on S897^22,28,35^, CK1 family kinases phosphorylate S901^28^ and PKC family kinases phosphorylate S892^29^. To identify the spectrum of kinases that phosphorylate the EphA2 kinase-SAM linker, we screened a collection of 298 kinases representing most of the serine/threonine kinase families in an *in vitro* kinase reaction with [γ-^33^P] ATP. As the substrate, we used a peptide corresponding to EphA2 residues D886–T908 (which include six residues before S892 and seven residues after S901). In this peptide S897 was already phosphorylated, to preclude phosphorylation by kinases targeting this site. Measurement of ^33^P incorporation revealed that multiple kinases readily phosphorylate the peptide substrate, with 15% of the kinases tested (i.e. 45 kinases) causing > 20,000 cpm ^33^P incorporation and accounting for ~70% of the total radioactivity incorporated (Fig. 6A; Table S2). Members of a number of different kinase families caused high peptide phosphorylation (> 50,000 cpm; Fig. 6B; Table S2). These include nearly all PKC family members, PAK3, members of the CK1 and NEK families, and a number of other kinases.

**Figure 6.**
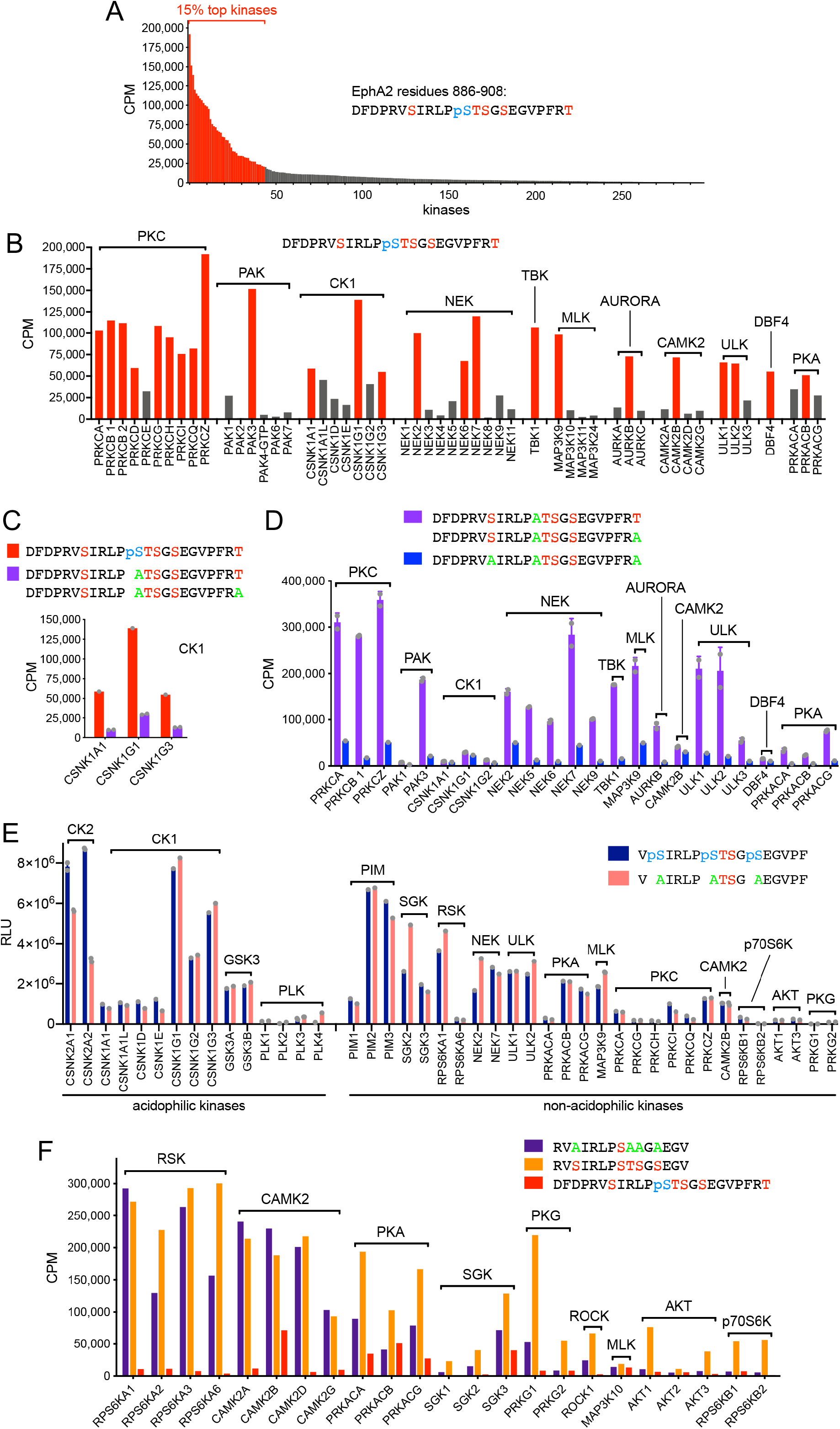
Multiple kinases phosphorylate the five serine/threonine residues in the EphA2 kinase-SAM linker. (**A**) Screen of 298 kinases using *in vitro* kinase reactions with [γ-^33^P] ATP and a peptide substrate containing S892, T898, S899, S901 and phosphorylated S897 identifies many kinases that phosphorylate residues other than S897. The 44 kinases (15% of those screened) mediating the highest ^33^P incorporation are shown in red. (**B**) The 24 kinases mediating the highest ^33^P incorporation (>50,000 CPM) are shown in red and other less active family members that were also tested are shown in grey. Kinase families are ordered based on the most active family member. (**C**) CK1 family members preferentially phosphorylate the peptide with phosphorylated S897 compared to two peptides with Ala replacing S897 and differing in the last residue (averaged together). (**D**) Comparison of two peptides containing S892 and differing in the last residue (averaged together) with a peptide in which S892 is replaced by Ala shows that many of the kinases identified in the screen mainly phosphorylate S892. Kinase families are ordered as in panel B. (**E**) Comparison of the phosphorylation of the two indicated peptides identifies kinases that can phosphorylate T898 and/or S899 in the presence or absence of prior S892, S897 and S901 phosphorylation. (**F**) ^33^P incorporation into peptides in which only S897 can be phosphorylated identifies kinase families that can phosphorylate this residue. Higher ^33^P incorporation into the peptide also containing the other four serine/threonine residues and residual ^33^P incorporation into the peptide with already phosphorylated S897 suggest that some of the kinases can also phosphorylate other residues. Error bars in C and D (for the measurements with two peptides differing only in the last residue) and in E (for duplicate measurements with CK2 kinases, PLK3, MAP3K9 and CAMK2B) represent SD. Individual data points in these panels are shown as gray dots. CPM, counts per minute measuring incorporated ^33^P; RLU, relative light units.

In the case of the constitutively active acidophilic CK1 family, we previously reported that these kinases phosphorylate EphA2 on S901 following phosphorylation of S897, which creates the necessary negatively charged motif^28^. This was confirmed in *in vitro* kinase assays with several CK1 family members, which did not efficiently phosphorylate peptide substrates in which the phosphorylated S897 was replaced by Ala (Fig. 6C). In addition, phosphorylation of peptides in which S892 was replaced by Ala revealed that most of the kinases identified in the screen preferentially phosphorylate S892 (Fig. 6D). Among these, we have previously shown that PKC family members play a particularly important role in S892 phosphorylation in cells^29^.

To identify kinases that phosphorylate T898 and/or S899, we used an ADP-Glo assay with two peptide substrates in which these two residues are the only possible phosphorylation sites (Fig. 6E). One of the peptides contains phosphorylated S892, S897 and S901, to identify kinases (such as acidophilic kinases^42–44^) that can generate a fourth and possibly a fifth phosphosite in the cluster. In the other peptide the three Ser residues were replaced by Ala, to identify kinases that can phosphorylate T898 and/or S899 in the absence of other phosphorylated residues. Among the acidophilic kinases (Fig. 6E left), we found that CK2 family kinases preferentially phosphorylate the peptide with three other phosphosites, suggesting that once S892, S897, and S901 are phosphorylated, the constitutively active CK2 can further increase the number of phosphosites. Notably, CK2 kinases optimally recognize substrates with a phosphosite or a negative charge at the + 3 position, such as T898 when S901 is phosphorylated and S899 (which is upstream of the negatively charged E902)^42,45^. The CK2 kinases were not among the most active kinases in the initial screen with the peptide phosphorylated only on S897 (Table S2). However, S897 phosphorylation may not be sufficient to promote phosphorylation of other cluster residues by CK2 (Fig. 6E). Some CK1 family members can also target T898 and/or S899, while the GSK3 family showed low activity (Fig. 6E).

Other (non-acidophilic) kinases were chosen for testing (Fig. 6E right) because of their residual activity towards the peptide containing T898, S899 and S901 as the only phosphorylatable residues (Fig. 6D). These kinases include NEK2 and NEK7, ULK1 and ULK2, MAP3K9, members of the PKC family, and CAMK2B (Fig. 6D). Additional kinases were chosen based on the prediction scores from the PhosphoNET Kinase Predictor online resource (phosphonet.ca), which takes into account only non-phosphorylated sequence motifs. The T898 motif has very low prediction scores, suggesting that this residue is unlikely to be phosphorylated when none of the other residues in the cluster is phosphorylated. The top predicted kinases for the S899 motif are those of the PIM, PKA, p70S6K, AKT, SGK, RSK and PKG families. Some of these kinases showed substantial phosphorylation of T898 and/or S899, with several of them preferring the peptide substrate lacking other phosphorylation sites (Fig. 6E). Taken together, these data suggest that T898, and possibly S899, can be phosphorylated by acidophilic kinases once one or more of the other residues in the cluster are phosphorylated. In addition, S899 and possibly T898 may also be phosphorylated by other kinases that do not require surrounding negative charges. Conceivably, once bound to the EphA2 kinase-SAM linker to phosphorylate S892 or S897, kinases such as PKC, RSK, AKT and PKA may processively phosphorylate additional residues^46–48^.

We also used a peptide containing only S897 as a potential phosphorylation site to screen a number of the kinases predicted by PhosphoNET to phosphorylate the S897 motif. This revealed new candidate S897 kinases, and most notably members of the CAMK2 family (Fig. 6F). Some of the kinases that phosphorylate the peptide containing only S897, show better activity towards a similar peptide that also contains the other four serine/threonine residues and/or still phosphorylate a peptide in which S897 is already phosphorylated. Thus, these kinases may also be able to phosphorylate other residues in the cluster.

The model emerging from these data is that a number of different kinases can phosphorylate the five serine/threonine residues in the EphA2 kinase-SAM linker, leading to concomitant phosphorylation of multiple residues that likely coordinately affect EphA2 signaling function by regulating the conformation of the EphA2 intracellular region.

## DISCUSSION

How the different parts of the intracellular region of Eph receptors – including the juxtamembrane segment, kinase domain, linker and SAM domain – are arranged and function together has remained an outstanding question, in part due to lack of structural information that extends beyond single domains and adjacent regions^38,39,49^. Our integrative structural biology characterization of most of the EphA2 intracellular region provides new insights into its conformation and dynamics. Our data suggest a model for the conformational dynamics that underlie EphA2 non-canonical signaling induced by cumulative serine/threonine phosphorylation of the kinase-SAM linker (Fig. S9A-B), extending previous findings on the role of tyrosine phosphorylation in Eph receptor canonical signaling^7–9,3^ (Fig. S9C). We uncovered two distinct molecular mechanisms that may function in concert to mediate the effects of linker phosphorylation through an orchestrated allosteric regulatory network. The first involves a shift in the equilibrium between a “closed” configuration of the EphA2 intracellular region (Fig. S9A) and an “open” more extended configuration (Fig. S9B) induced by the accumulation of phosphorylation sites in the linker. The second involves allosteric rearrangements in the kinase domain and juxtamembrane segment induced by phosphorylation of S892 and, to a lesser extent, S897, suggesting a link between canononical and non-canonical signaling (Fig. S9B,C).

Our hydrogen-deuterium exchange data suggest that in the absence of linker phosphorylation the electronegative surface of the EphA2 kinase domain interacts with the electropositive surface of the SAM domain favoring the closed conformation (Fig. S9A). In addition, molecular dynamics simulations have previously suggested that in the absence of tyrosine phosphorylation electropositive patches on the opposite side of the kinase domain interact with anionic phospholipids on the cytosolic side of the plasma membrane and that these interactions are facilitated by positive charges in the membrane-proximal portion of the juxtamembrane segment, which also interact with phospholipids^34,50^(Fig. S9A). Furthermore, the membrane-distal portion of the juxtamembrane segment interacts with the kinase domain stabilizing the catalytically inactive state. Thus, the inactive form of EphA2 lacking both canonical and non-canonical signaling is in a closed, compact configuration closely associated with the plasma membrane (Fig. S9A).

We propose that serine/threonine phosphorylation of the kinase-SAM linker promotes an allosteric rearrangement of the EphA2 intracellular region, shifting the equilibrium to an open conformation in which the interaction between kinase and SAM domains is weakened or disrupted (Fig. S9B). EphA2 non-canonical signaling involving S897 phosphorylation was discovered more than ten years ago^22^. Although S897 is part of a cluster of five phosphorylated residues, prior studies have focused essentially only on this serine^26^. The importance of S897 phosphorylation is highlighted by the effects of the S897A mutation, which prevents phosphorylation and drastically impairs the ability of EphA2 to mediate non-canonical signaling^22,28,35^. Our findings, however, suggest that the role of S897 phosphorylation does not depend only on the modification of this single residue but also involves promoting phosphorylation of other linker residues. Therefore, we hypothesized that EphA2 non-canonical signaling is regulated by concomitant phosphorylation of multiple residues in the EphA2 kinase-SAM linker. Interestingly, four of the five phosphorylation sites are conserved in another Eph receptor, EphA1, supporting the functional importance of the phosphorylation cluster as well as the ability of EphA1 to also mediate non-canonical signaling. In fact, evidence shows that serine/threonine phosphorylation sites tend to cluster in disordered regions of proteins, where they are coordinately regulated and functionally cooperate^40,48,51^. Phosphosites in a cluster can accumulate until a response threshold is reached, enable gradual effects, and/or potentiate the robustness of phosphorylation-dependent responses.

Several mechanisms could lead to the observed concomitant phosphorylation of multiple residues in the EphA2 linker. First, phosphorylation of some linker residues can promote phosphorylation of additional residues by acidophilic kinases^42,47,52^. Indeed, we found that CK1 and CK2 preferentially phosphorylate the EphA2 linker peptide when other residues are already phosphorylated (Fig. 6C,E)^28^. A second mechanism likely involves processive phosphorylation^46–48^. Once bound to EphA2 in order to phosphorylate a highly compatible consensus motif, a kinase can continue to phosphorylate multiple nearby residues even if they are part of less optimal motifs. In agreement with this, basophilic kinases such as PKA, AKT, and RSK are predicted to phosphorylate not only S897 but also two other linker residues (S892 and S899; phosphonet.ca). Perhaps also consistent with this, we have previously found that PKA activation increases EphA2 linker phosphorylation on multiple residues^28^ and we show here that PKA and other kinases are able to phosphorylate multiple linker residues (Fig. 6). A third mechanism is the coincident phosphorylation of different residues by kinases that are concomitantly regulated, for example by oncogenic transformation or because they are part of the same signaling network^29,53–55^. All three mechanisms conceivably contribute to the cumulative phosphorylation of the EphA2 kinase-SAM linker. Further supporting the notion that the EphA2 linker can be highly phosphorylated, we found that a number of kinases can add a fourth and possibly a fifth phosphorylation site to the cluster in *in vitro* kinase reactions (Fig. 6E). Taken together, these findings imply that regulation of EphA2 phosphorylation leading to non-canonical signaling is more complex than previously envisioned. Accordingly, we have identified multiple kinases that can phosphorylate each serine/threonine residue in the EphA2 linker in *in vitro* kinase reactions (Table S2). While a number of these kinases have already been shown to phosphorylate EphA2 in cells^22,26,28,29,35^, additional studies are needed to further define the repertoire of kinases that phosphorylate the EphA2 kinase-SAM linker in cells to regulate non-canonical signaling in response to different stimuli.

A key question is how phosphorylation of the EphA2 linker regulates non-canonical signaling in the cell. Linkers are known to play a crucial role in allosteric propagation of information between protein modules by communicating conformational and/or dynamic alterations^56–59^. Due to their flexible nature, they can serve as hinges, enabling reorientation of protein domains with respect to each other to regulate protein function. Length, amino acid composition and posttranslational modifications such as phosphorylation can profoundly impact the conformation and dynamics of linkers, affecting their allosteric behavior. By using hydrogen-deuterium exchange, we uncovered cumulative effects of the negative charges mimicking phosphorylation of the linker. We propose, that these negative charges cause a progressive allosterically driven reorientation of the EphA2 intracellular region shifting the equilibrium from the closed conformation, in which electrostatically complementary portions of the kinase and SAM domains are protected (and thus presumably interact with each other), to a more extended open conformation (Fig. S9A,B). The model that the linker is flexible, and its phosphorylation regulates the configuration of the EphA2 intracellular region, is supported by the variation in the relative arrangement of the two domains observed in our crystal structures. In addition, we found that the E820K-E825K double mutation, which disrupts the electronegative surface of the EphA2 kinase domain, and thus likely its interaction with the SAM domain, has similar effects on hydrogen-deuterium exchange as multiple phosphomimetic mutations in the kinase-SAM linker. Interestingly, we also found that in the EphA2 open conformation, the linker is less solvent exposed, suggesting that phosphorylation of the linker may promote its interaction with other parts of EphA2 that perhaps further stabilize the open conformation. In summary, we speculate that the cumulative effects of different serine/threonine phosphorylation sites in the linker represent a barcode that allows fine-tuning of non-canonical signaling strength by progressively shifting the EphA2 conformational equilibrium towards the open conformation.

The EphA2 open conformation promoted by linker phosphorylation could expose kinase and SAM domain surfaces not readily accessible in the closed conformation for the binding of effector proteins that mediate non-canonical signaling (Fig. S9B). The association best known to be important for EphA2 non-canonical signaling is that involving the RHOG guanine nucleotide exchange factor Ephexin4 (gene name ARHGEF16), which leads to downstream activation of the RHOG-RAC and PI3K-AKT axes to promote cell migration, survival and proliferation^60–62^. Ephexin4 has been reported to interact with the EphA2 kinase domain in co-immunoprecipitation experiments^60,63^. However, it remains to be determined whether the EphA2 association with Ephexin4 is direct or depends on an intermediary protein. Favoring the latter, in size-exclusion chromatography, isothermal titration calorimetry and recombinant co-expression/co-purification experiments, we did not detect binding between the purified Ephexin4 DH-PH domains and the EphA2 intracellular region with or without the kinase domain or the S897E/S901E phosphomimetic mutations (unpublished data). Furthermore, although we confirmed weak EphA2-Ephexin4 association in pulldowns from transiently transfected HEK293 cells, we did not observe effects of the EphA2 S897A mutation or SAM domain deletion on the interaction (unpublished data). The SRC family kinase LYN has been recently found to preferentially interact with EphA2 phosphorylated on S897 under conditions of high extracellular matrix rigidity, triggering a mechanotransduction pathway that drives breast cancer invasion and metastasis^25^. Furthermore, the phosphomimetic S897E mutation has been recently shown to promote EphA2 association with the RHO family GTPase CDC42 leading to ameboid movement, transendothelial migration and metastasis of BRAF inhibitor-resistant melanoma cells^64^. The importance of other linker phosphorylation sites in these effects of EphA2 non-canonical signaling remains to be investigated. It is also not known whether phosphorylated EphA2 linker residues may modulate association with binding partners through other mechanisms, such as by creating specific binding motifs or introducing negative charges.

Our FRET data suggest that the phosphorylation state of the kinase-SAM linker also plays a minor role in regulating lateral EphA2 interactions on the cell surface. We found that mutations abolishing phosphorylation sites in the linker modestly decrease EphA2 dimerization in the absence of ephrin stimulation as compared to phosphomimetic mutations or phosphorylated residues in EphA2 WT. The most notable effect on EphA2 dimerization was observed when all five phosphosites in the kinase-SAM linker are mutated to alanine. Thus, intramolecular interaction of the SAM domain with the kinase domain in the closed configuration may impair dimerization, perhaps by preventing intermolecular interactions between the positively and negatively charged surfaces on opposite sides of the EphA2 kinase domain that can occur more readily in the open configuration. On the other hand, the negative charges in the kinase-SAM linker may cause repulsion between EphA2 molecules positioned side-by-side in the plasma membrane, thus moderating the positive effects of linker phosphorylation on dimerization.

The structural information we obtained also suggests that E820 in the αFG loop of the kinase domain plays a central role as a sensor for linker phosphorylation in an allosteric network connecting the SAM domain, linker, kinase domain and juxtamembrane segment. In particular, E820 appears to be a sensor for S892 phosphorylation and, to a lesser extent, S897 phosphorylation. Electrostatic repulsion between E820 and phosphorylated S892 or S897 could at least in part explain why the region around E820 becomes more solvent exposed in the EphA2 S892E mutant and to a lesser extent in the S897E mutant in HDX experiments. Importantly, E820 is in a different conformation in the EphA2 WT structure compared to the EphA2 S897E/S901E structure (Fig. S10), supporting this hypothesis. Interestingly, in an EphA3 structure lacking the SAM domain^38^, the equivalent E827 residue points even further upwards and away from the linker (Fig. S10). The EphA3 linker is not known to be phosphorylated, but includes the negatively charged D904 at the position corresponding to S897^38^. An allosteric linkage of the EphA3 kinase-SAM linker with the αFG loop, activation loop and juxtamembrane segment mediated by E827 and W826 (corresponding to EphA2 E820 and W819) has previously been proposed^39^. Similarly, in EphA2 changes in E820 and the αFG loop may be relayed to the activation loop (which is located just above E820; Fig. S10), the juxtamembrane segment, and the SAM domain (which interacts with the negatively charged surface formed by the αFG loop and adjacent regions). These observations suggest that the E820K mutation has multiple effects on the configuration of the E820K/E825K mutant intracellular region, not only impeding binding of the SAM domain to the negatively charged surface of the kinase domain but also disrupting the allosteric network involving the kinase-SAM linker.

We have examined the structure and dynamics of the EphA2 intracellular region in the absence of tyrosine phosphorylation. In other Eph receptors, phosphorylation of one or both conserved tyrosines in the juxtamembrane segment disrupts intramolecular inhibitory interactions, enabling receptor kinase activity and canonical signaling^7–9,38^. The different interaction of Y594 in our crystal structures and in other EphA2 structures suggests that the juxtamembrane segment has a dynamic conformation even in the absence of phosphorylation. Furthermore, dissociation of the EphA3 juxtamembrane segment from the kinase domain due to tyrosine phosphorylation has been shown to weaken interaction of the kinase domain with the plasma membrane, resulting in a more elongated conformation of the intracellular region that extends further away from the plasma membrane and has distinctive functional properties^65^(Fig. S9C). Our HDX-MS experiments identified allosteric coupling of the kinase-SAM-linker, activation loop and the juxtamembrane segment, suggesting a concerted interrelationship of the different parts of the EphA2 intracellular region. Notably, *in vitro* kinase reactions with the recombinant EphA3 intracellular region lacking the SAM domain suggest that the same allosteric network is involved in the regulation of receptor autophosphorylation on tyrosine residues^39^, implying that canonical and non-canonical signaling may influence each other (Fig. S9, black double-headed arrow). It will be important in future experiments to investigate the interplay of kinase-SAM linker serine/threonine phosphorylation and autophosphorylation on tyrosine residues in EphA2 signaling and function. In addition, previous data show that in certain cellular contexts EphA2 canonical signaling can downregulate non-canonical signaling through inhibition of ERK-RSK and AKT^20,22,28^(Fig. S9, blue inhibitory arrow).

In conclusion, we have shown that the EphA2 kinase-SAM linker can be simultaneously phosphorylated on multiple serine/threonine residues and have identified a repertoire of kinases able to phosphorylate these residues. Furthermore, we have uncovered how cumulative phosphorylation on multiple linker residues can allosterically affect the configuration of the EphA2 intracellular region. Since EphA2 kinase-SAM linker phosphorylation has been widely implicated in cancer malignancy, our findings can inform further mechanistic studies of EphA2 function as well as therapeutic strategies to target EphA2 signaling. Our studies also have implications for the regulation of other members of the Eph receptor family.

## METHODS

### Antibodies

Antibodies recognizing EphA2 (rabbit mAb #6997) and the phosphorylated S897 motif (rabbit mAb #6347) were purchased from Cell Signaling Technology. Affinity purified rabbit antibodies recognizing the phosphorylated S892 or S901 motifs were generated and validated as described^28,29^.

### Cell lines

The following cell lines were purchased from the American Type Culture Collection (Manassas, VA): PC3 androgen-independent prostate cancer (CRL-1435), A549, Calu-3 and H1648 lung adenocarcinoma (CCL-185, HTB-55 and CRL-5882 respectively), HCC1937 triple negative breast cancer (CRL-2336), SW626 colon adenocarcinoma (HTB-78), SKOV3 ovarian serous cystadenocarcinoma (HTB-77) and HEK 293T human embryonic kidney (CRL-3216). The MEL-JUSO melanoma cell line was purchased from the DSMZ (ACC 74); the HOP62 lung adenocarcinoma cell line from the tumor/cell line repository of the Developmental Therapeutics Program, Division of Cancer Treatment and Diagnosis, National Cancer Institute; and the HEK 293AD human embryonic kidney cell line from Cell Biolabs, Inc (AD-100). The BxPC3 pancreatic cancer cell line was kindly provided by P. Itkin-Ansari (Sanford Burnham Prebys Medical Discovery Institute), the PANC1 cell line by F. Levine (Sanford Burnham Prebys Medical Discovery Institute); the U87, T98G and U251-MG glioblastoma cell lines by W. Stallcup (Sanford Burnham Prebys Medical Discovery Institute); the BT549 breast cancer cell line by R. Maki (Sanford Burnham Prebys Medical Discovery Institute); the MDA-MB-468 breast cancer cell line by K. Vuori (Sanford Burnham Prebys Medical Discovery Institute); and the MDA-MB-231 breast cancer cell line by J. Smith (Sanford Burnham Prebys Medical Discovery Institute). The MEL-JUSO cell line was stably infected with the pLVX-IRES-Neo lentivirus encoding EphA2 wild-type with an N-terminal FLAG tag^28^.

The PC3, MDA-MB-231, BT549, H2009, PANC-1, T98G, U87 and U251-MG cell lines were authenticated by performing short tandem repeat analysis on isolated genomic DNA with the GenePrint® 10 System (Promega), and peaks were analyzed using GeneMarker HID (Softgenetics). Allele calls were searched against short tandem repeat databases^66^.

A549, BxPC3, H1648, HCC1937, HOP62, PC3 and MEL-JUSO cells were cultured in RPMI 1640 medium (Gibco); BT549, Calu-3, T98G, MDA-MB-231, MDA-MB-468, PANC1, SW626, U251-MG and U87 cells were cultured in Dulbecco’s Modified Eagle Medium (DMEM; Corning) and SKOV3 cells were cultured in McCoy’s containing 10% FBS. All culture media contained 10% fetal bovine serum, antimycotics and antibiotics. HEK 293T cells used in FRET experiments were cultured in DMEM supplemented with 10% fetal bovine serum, 3.5 g l^−1^D-glucose, and 1.5 g l^−1^sodium bicarbonate.

### Kinase reactions

The peptide substrates were synthesized by Kinexus (Vancouver, Canada) at >95% or > 98% purity. The recombinant protein kinases were cloned, expressed and purified using Kinexus proprietary methods and underwent routine quality control testing. The assay conditions were optimized to yield acceptable enzymatic activity. In addition, the assays were optimized to give high signal-to-noise ratio.

Protein kinase assays were performed by Kinexus, in most cases using a radioisotope assay format. These assays were performed in a final volume of 25 μl, including 5 μl diluted active kinase (~10-50 nM final concentration), 5 μl stock peptide substrate solution (500 μM final concentration), 10 μl kinase assay buffer, 5 μl [γ-^33^P]ATP (250 μM stock solution, 0.8 μCi). The assay was initiated by the addition of [γ-^33^P]ATP and the reaction mixture incubated at room temperature for 30 min. The assay was terminated by spotting 10 μl of the reaction mixture onto a multiscreen phosphocellulose P81 plate, which was then washed 3 x 15 min in a 1% phosphoric acid solution. The radioactivity on the P81 plate was measured in the presence of scintillation fluid in a Trilux scintillation counter.

Some protein kinase assays were performed using the ADP-Glo^™^ assay kit (Promega), which measures the generation of ADP in a kinase reaction. Generation of ADP leads to an increase in luminescence signal in the presence of ADP-Glo^™^. The protein kinase assays were performed at 30°C for 30 min in a final volume of 25 μl, including 5 μl diluted active protein kinase to obtain a previously determined optimal concentration, 5 μl of a 125 μM stock peptide substrate solution (25 μM final concentration), 5 μl kinase assay buffer, 5 μl 10% DMSO, 5 μl ATP stock solution. The assay was started by incubating the reaction mixture in a 96-well plate at room temperature for 30 min. The assay was terminated by the addition of 25 μl of ADP-Glo^™^ Reagent (Promega). The 96-well plate was shaken and then incubated for 40 min at room temperature. Then, 50 μl of Kinase Detection Reagent was added, the 96-well plate was shaken and then incubated for further 30 min at room temperature. The 96-well reaction plate was read using the ADP-Glo^™^ Luminescence Protocol on a GloMax plate reader (Promega; Cat# E7031).

### Immunoprecipitation and immunoblotting

Cells were collected when they reached ~90% confluency. For immunoprecipitation, cells were collected in ice-cold modified RIPA buffer (50 mM Tris-HCl pH7.6, 150 mM NaCl, 1% Triton X-100, 0.5% sodium deoxycholate, 0.1% SDS, and 2 mM EDTA) supplemented with Halt Protease and Phosphatase Inhibitor cocktail (PI78442; Thermo Fisher Scientific). Lysates were centrifuged at 13,000 g for 10 min at 4°C to remove insoluble material. Supernatants were pre-cleared by incubation with Sepharose beads (4B200; Sigma Aldrich) for 15 min at 4°C on a rotator. Each immunoprecipitation was performed using 6 μg EphA2 antibody (05-480; EMD Millipore) coupled to 20 μl GammaBind Plus sepharose beads (17-0886-02; GE Healthcare) for 2 hours at 4°C. Immunoprecipitates were washed three times with ice-cold modified RIPA buffer and once with phosphate buffered saline (PBS), and eluted by heating at 95°C for 2 min in SDS-PAGE sample buffer. For immunoblotting, cells were collected in SDS-PAGE sample buffer and lysates were heated at 95°C for 2 min followed by a brief sonication. After SDS-PAGE and transfer to a PVDF membrane, blots were incubated for 1 hour in blocking buffer (0.1% Tween-20, 5% BSA in PBS) and then overnight at 4°C with primary antibodies diluted in blocking buffer. Membranes were then incubated for 1 hour at room temperature with goat-anti-rabbit secondary antibodies conjugated to HRP (A16110; Invitrogen/ThermoFisher Scientific) followed by ECL chemiluminescence detection (RNP2106; GE Healthcare Bio-Sciences, Piscataway, NJ) using a ChemiDoc gel imaging system (Bio-Rad).

### Recombinant EphA2 expression and purification

The cDNA sequence coding for the intracellular portion of human EphA2 (residues 590–976) was cloned into the pETNKI-LIC vector^67^, which encodes an N-terminal 6xHis-tag followed by a 3C protease cleavage site in a pET29 vector backbone. Mutations were introduced in constructs using standard site-directed mutagenesis. The sequences of all PCR-amplified portions of constructs were verified by Sanger sequencing (Retrogen, San Diego).

EphA2 was expressed in *E. coli* BL21(DE3) grown in 2xYT medium (BD Difco) at 20°C overnight and purified using Ni-NTA agarose (Qiagen) followed by cleavage of the His-tag with 3C protease and dephosphorylation with bovine alkaline phosphatase (Sigma). Further purification was performed by ion exchange chromatography (Source Q for WT and the S897E, S897E/S901E, and E820K/E825K mutants and Source S for the 3E and 5E mutants) equilibrated in 10 mM NaCl, 10 mM HEPES pH 7.9 and eluted with a linear NaCl gradient up to 500 mM. EphA2 was concentrated to 5-7 mg ml^−1^. For HDX-MS, the cDNA sequence coding for the intracellular portion of human EphA2 (residues S570-D976) was cloned into the pET29a(+) vector (Novagen) to express EphA2 protein with a C-terminal 6xHis-tag. The protein was purified as described above, but no 3C protease cleavage was performed and the protein was further purified on a Superdex 200 Increase 10/300 GL size-exclusion chromatography column (GE Healthcare) equilibrated in 100 mM NaCl, 10 mM HEPES pH 7.9. All proteins were flash frozen in ~100 μl aliquots in liquid nitrogen and stored at −80°C.

### Crystallization and structure solution

For crystallization, EphA2 (residues D590-I976, ~7 mg ml^−1^) was mixed with 1.7 mM AMPPcP and 10 mM MgCl_2_, and initial crystals were obtained with the Hampton Index HT screen at 14°C. Crystals were optimized by increasing drop size and varying protein to precipitant ratio. Crystals of the S897E/S901E mutant (~5 mg ml^−1^) were additionally optimized using the Hampton Additive Screen HT. Final crystallization conditions are listed in Table S1. Crystals were cryoprotected by step-wise transfer to reservoir solutions supplemented with 5-15% glycerol and cryocooled in a nitrogen stream at 100 K. Diffraction data were collected on a rotating anode X-ray generator (Rigaku FR-E superbright) at 100 K and processed in XDS^68^ or MOSFLM^69^ and with software from the CCP4 suite^70^. Phases were obtained using molecular replacement in Phaser^71^ with the EphA2 kinase domain from PDBID 4PDO (unpublished) and the EphA2 SAM domain from PDBID 3KKA (unpublished) as search models. Model building and refinement were respectively performed in Coot^72^ and Refmac^73^ or Phenix^74^. The final model was validated using MolProbity^75^. Data collection and refinement statistics are reported in Table S1. Crystallographic models and data have been deposited in the PDB under accession numbers 7KJA (EphA2 WT residues 590-976), 7KJB (EphA2 S897E/S901E residues 590-976) and 7KJC (EphA2 S901E residues 590-976).

### Size exclusion chromatography–small-angle X-ray scattering (SEC-SAXS)

EphA2 proteins were dialyzed into 100 mM NaCl, 10 mM HEPES pH 7.9 and their concentration was adjusted to 2 mg ml^−1^. SAXS data were collected at the Argonne National Laboratory Advanced Photon Source Biocat SAXS/WAXS beamline 18-ID, using an inline size-exclusion chromatography set-up (AKTA pure with Superdex 200 10/300 GL column; GE Healthcare). Two-dimensional scattering data were radially averaged, normalized to sample transmission, and scatter patterns from the apex of the size-exclusion chromatography elution peak were averaged. Background scatter (scatter patterns from early in the SEC run) was subtracted. The ATSAS suite of software^76^ and RAW 1.5.2^77^ were used for data analysis. These included: (i) Guinier analysis (PRIMUS^78^) to calculate the pair-wise intra-atomic distance distribution function *P*(*r*) and maximum particle dimension and (ii) *D*_max_(GNOM_79_) to fit and refine structural models to experimental SAXS data (CRYSOL and SREFLEX) and for bead model generation (DAMMIF, DAMAVER, and DAMMIN). Bead models and crystal structures were superimposed using the ATSAS subcomb script. All structural models were illustrated using PyMOL.

### Hydrogen-deuterium exchange–mass spectrometry (HDX-MS)

HDX-MS was performed at the Biomolecular and Proteomics Mass Spectrometry Facility (BPMSF) of the University California San Diego, using a Synapt G2Si system with HDX technology (Waters Corporation) according to methods previously described^80^. Briefly, deuterium exchange reactions were performed using a Leap HDX PAL autosampler (Leap Technologies, Carrboro, NC). D_2_O buffer was prepared by lyophilizing 10 mM HEPES pH 7.9, 100 mM NaCl initially dissolved in ultrapure water and redissolving the powder in the same volume of 99.96% D_2_O (Cambridge Isotope Laboratories, Inc., Andover, MA) immediately before use. Deuterium exchange was measured in triplicate at each time point (0 min, 0.5 min, 1 min, 2 min, 5 min). For each deuteration time point, 4 μl of protein (5 μM) was held at 25 °C for 5 min before being mixed with 56 μl of D_2_O buffer. The deuterium exchange was quenched for 1 min at 1°C by combining 50 μl of the deuteration reaction with 50 μl of 250 mM tris(2-carboxyethyl)phosphine (TCEP) pH 2.5. The quenched sample (90 μl) was then injected in a 100 μl sample loop, followed by digestion on an in-line pepsin column (Immobilized Pepsin, Pierce) at 15°C. The resulting peptides were captured on a BEH C18 Vanguard precolumn, separated by analytical chromatography (Acquity UPLC BEH C18, 1.7 μm 1.0 × 50 mm, Waters Corporation) using a 7–85% acetonitrile gradient in 0.1% formic acid over 7.5 min, and electrosprayed into the Waters Synapt G2Si quadrupole time-of-flight mass spectrometer. The mass spectrometer was set to collect data in the Mobility, ESI + mode; mass acquisition range of 200– 2000 (m/z); scan time 0.4 s. Continuous lock mass correction was accomplished with infusion of leu-enkephalin (m/z = 556.277) every 30 s (mass accuracy of 1 ppm for calibration standard).

For peptide identification, the mass spectrometer was set to collect data in mobility-enhanced data-independent acquisition (MS^E^), mobility ESI + mode instead. Peptide masses were identified from triplicate analyses and data were analyzed using the ProteinLynx global server (PLGS) version 2.5 (Waters Corporation). Peptide masses were identified using a minimum number of 250 ion counts for low energy peptides and 50 ion counts for their fragment ions; the peptides also had to be larger than 1,500 Da. The following cutoffs were used to filter peptide sequence matches: minimum products per amino acid of 0.2, minimum score of 7, maximum MH + error of 5 ppm, and a retention time RSD of 5%. In addition, the peptides had to be present in two of the three ID runs collected.

The peptides identified in PLGS were then analyzed using DynamX 3.0 data analysis software (Waters Corporation). Peptides containing mutated residues were manually assigned. The relative deuterium uptake for each peptide was calculated by comparing the centroids of the mass envelopes of the deuterated samples with the undeuterated controls following previously published methods^81^. The peptides reported on the coverage maps are actually those from which data were obtained. For all HDX-MS data, 3 technical replicates were analyzed. The deuterium uptake was corrected for back-exchange using a global back exchange correction factor (typically ~25%) determined from the average percent exchange measured in disordered termini of various proteins^82^. Deuterium uptake plots in Fig S5 and coverage maps in Fig S6 were generated in DECA (github.com/komiveslab/DECA) and the data in Fig S5 are fitted with an exponential curve for ease of viewing^83^.

### Analytical ultracentrifugation

Sedimentation velocity experiments were performed using a ProteomeLab XL-I (Beckman Coulter) analytical ultracentrifuge. EphA2 590-976 sample with OD_280nm_ = 0.5 (~9 μM) in 10 mM HEPES pH 7.9, 100 mM NaCl was loaded into 2-channel cells and spun in an An-50 Ti 8-place rotor at 40,000 rpm and 20°C for 15 hours. Data were analysed using Sedfit software (P. Schuck, NIH/NIBIB)^84^.

### FRET imaging and analysis

HEK 293T cells were seeded at a density of 2×10^5^ cells/dish in 35 mm glass bottom collagen-coated dishes (MatTek Corporation) and cultured for 24 hours at 37 °C in 5% CO_2_. The cells were transiently co-transfected with EphA2-mTurquoise and EphA2-EYFP WT and 5E, 3E, S897E, S892E, 5A, ASAAA, S897A, or S892A mutants in pcDNA3 using the Lipofectamine 3000 reagent (ThermoFisher Scientific/Invitrogen). Cells were transfected with 0.5-2 μg total DNA at a 1:2 EphA2-mTurquoise:EphA2-EYFP ratio. Twelve hours later, the cells were washed twice with starvation medium to remove traces of phenol red, and serum-starved overnight to ensure no soluble ligands were present.

FRET experiments were performed following published protocols^41,85,86^. HEK 293T cells expressing full-length EphA2 WT and mutants were imaged under reversible hypo-osmotic conditions. For this, the medium was replaced with hypotonic swelling medium (25 mM HEPES in serum free medium) about 5 min prior to imaging. The cells were then imaged for approximately 1 hour. Cell osmotic swelling is necessary to eliminate the numerous “wrinkles” present in the plasma membrane of live cells, which hinder measurement of receptor concentration in the plasma membrane (see below)^87^.

A two-photon microscope equipped with the OptiMis True Line Spectral Imaging system (Aurora Spectral Technologies) was used to image the cells, and a MaiTai laser (Spectra-Physics) was used to generate femtosecond mode-locked pulses^88,89^. Two images were acquired for each cell: one upon excitation at 840 nm to mainly excite the donor, and a second one at 960 nm to primarily excite the acceptor^41^. The FSI-FRET method was used to measure the FRET efficiency and the concentrations of donor (EphA2-mTurquoise) and acceptor (EphA2-EYFP) in small cell plasma membrane areas. Calibration solutions of purified soluble EYFP and mTurquoise, produced using a published protocol^90^, were used to convert pixel-level intensities of the images into EphA2 concentrations as previously described^41^.

The FRET efficiency due to EphA2 dimerization depends on the fraction of receptors that are dimeric, *f_D_*, and on the acceptor fraction, *x_A_*, according to

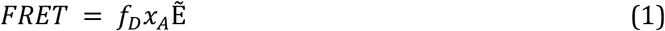

The Ẽ in equation (1) is a constant called the “intrinsic FRET”, which represents the FRET efficiency in a dimer containing a donor molecule and an acceptor molecule^91,92^. Intrinsic FRET is a structural parameter that depends only on the separation and orientation of the two fluorescent proteins in the dimer, and not on the dimerization propensity. The distance between the fluorescent proteins in the dimer, *d*, under the assumption of free fluorescent protein rotation, is given by^91^:

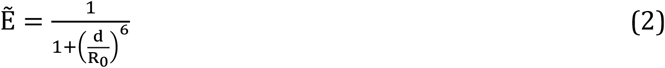

where *R_0_* is 54.5 Å, the Förster radius for the mTurquoise-EYFP FRET pair.

The following equation, which provides a link between the experimentally derived parameters *FRET*, *x_A_* and the total EphA2 concentration [*T*] (including the concentrations of both EphA2-mTurquoise and EphA2-EYFP), and the two unknowns, *K_diss_* and Ẽ, was used to fit the data as previously described^41^.

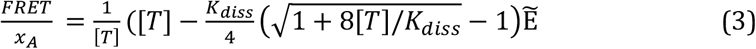

## DATA AVAILABILITY

Atomic structures and diffraction data have been deposited in the PDB with accession codes 7KJA (EphA2 WT), 7KJB (EphA2 S897E/S901E) and 7KJC (EphA2 S901E). HDX-MS data have been deposited in the Mass Spectrometry Interactive Virtual Environment (MassIVE, massive.ucsd.edu) with accession code MSV000086658. Kinase screening data are presented in Supplementary Table S2. All other data supporting the conclusions are available with the article. Raw data, including immunoblots, are available upon request to the authors.

## ACKNOWLEDGMENTS

The authors thank P. Hornbeck (PhosphoSitePlus) for providing original data on phosphosites in individual peptides from the PhosphoSite database, Alejandro Conde-Perez for generating the MEL-JUSO-EphA2 stable cell line, B. Emerling for providing the SW626 cell line, P. Itkin-Ansari for the BxPC3 cell line, F. Levine for the PANC1 pancreatic cancer cell line, K. Vuori for the SKOV3 and MDA-MB-468 cell lines, J. Smith for the MDA-MB-231 cell line, R. Maki for the BT549 cell line, W. Stallcup for the U87, T98 and U251-MG cell lines, the staff of the BioCAT Advanced SAXS Training Course, in particular W. Shang, for access to the SAXS beamline and help with SAXS data analysis, J. Murphy (WEHI) for discussing SAXS data analysis, A. Bobkov for collecting and analyzing AUC data, S. Silletti (UCSD Biomolecular and Proteomics Mass Spectrometry (BPSM) Facility) for HDX-MS data collection and analysis and Ryan Lumpkin for help with DECA. This work was supported by NIH grant GM131374 and institutional funds to EBP, and National Cancer Institute Cancer Center Support grant P30 CA030199, which supported institutional Core Facilities and funds for a pilot project. The UCSD BPSM Facility is supported by the NIH shared instrumentation grant number S10 OD016234 (Synapt-HDX-MS). This project also used resources of the Advanced Photon Source, a U.S. Department of Energy (DOE) Office of Science User Facility operated for the DOE Office of Science by Argonne National Laboratory under Contract No. DE-AC02-06CH11357. The work for this project was supported by grant 9 P41 GM103622 from the National Institute of General Medical Sciences of the National Institutes of Health. Use of the BioCAT Pilatus 3 1M detector was provided by grant 1S10OD018090-01 from NIGMS. The content is solely the responsibility of the authors and does not necessarily reflect the official views of the National Institute of General Medical Sciences or the National Institutes of Health.

## AUTHOR CONTRIBUTIONS

B.C.L. and E.B.P. designed the overall study and interpreted the results; T.P.L. and K.H. designed and interpreted the FRET experiments; B.C.L., M.P.G., T.P.L., and M.W.M. performed experiments; B.C.L. and E.B.P. wrote the manuscript with input from T.P.L. and K.H.; E.B.P. and K.H. secured funding.

