## Supplemental Tables & Figures for "Allosteric Regulation of the EphA2 Receptor Intracellular Region by Serine/Threonine Kinases"

**Table S1. Crystallographic data collection and refinement statistics**

| | EphA2 WT | EphA2 S897E/S901E | EphA2 S901E |
|---|---|---|---|
| **Crystal** | | | |
| PDB ID | 7KJA | 7KJB | 7KJC |
| Space group | I2 | $P3_221$ | I2 |
| Cell dimensions | | | |
| a, b, c (Å) | 123.1, 55.0, 135.7 | 94.5, 94.5, 99.7 | 121.6, 54.6, 135.5 |
| α, β, γ (°) | 90, 94.86, 90 | 90, 90, 120 | 90, 95.2, 90 |
| Crystallization conditions | 0.2 M $MgCl_2$, 0.1 M Bis-Tris pH5.5, 25% PEG 3,350 | 4.5% Tacsimate, 0.09 M HEPES pH 7.0, 9% PEG-MME 5K, 0.1 M CsCl | 0.2 M $MgCl_2$, 0.1 M Bis-Tris pH5.5, 25% PEG 3,350 |
| **Data processing statistics** | | | |
| Resolution (Å) | 20.49 - 1.75 (1.78-1.75) | 29.53 - 2.80 (2.95 - 2.80) | 29.25 - 2.30 (2.38-2.30) |
| $R_{merge}$ | 0.059 (0.690) | 0.285 (1.956) | 0.089 (0.656) |
| Reflections | 327758 (15791) | 142325 (20517) | 146921 (14220) |
| Unique Reflections | 91439 (4522) | 13051 (1871) | 39208 (3887) |
| $\langle I/\sigma I \rangle$ | 13.4 (1.4) | 10.2 (1.4) | 12.0 (1.8) |
| $CC_{1/2}$ | 0.997 (0.599) | 0.992 (0.554) | 0.995 (0.643) |
| Completeness (%) | 99.9 (100) | 99.9 (100.0) | 99.8 (99.8) |
| Redundancy | 3.6 (3.5) | 10.9 (11.0) | 3.7 (3.7) |
| **Model** | | | |
| EphA2/peptide complexes per asu | 2 | 1 | 2 |
| No. atoms (non-H) | 6472 | 2913 | 6277 |
| EphA2 | 5764 | 2882 | 5847 |
| AMPPcP | 62 | --- | 62 |
| Water | 615 | 30 | 343 |
| Other solvent | 31 | 1 | 25 |
| Average B-factors (non-H) | | | |
| EphA2 | 32.7 | 57.3 | 43.2 |
| AMPPcP | 41.8 | --- | 51.6 |
| Water | 34.6 | 36.9 | 37.8 |
| Other solvent | 45.5 | 121.1 | 55 |
| **Refinement statistics** | | | |
| Resolution (Å) | 20.48 - 1.75 (1.81 - 1.75) | 29.53 - 2.8 (2.9 - 2.8) | 29.25 - 2.3 (2.38 - 2.3) |
| No. reflections | 90866 (9053) | 13027 (1269) | 39703 (3948) |
| $R_{work}/R_{free}$ | 0.1883 (0.2717) / 0.2173 (0.3019) | 0.1976 (0.2849) / 0.2461 (0.3097) | 0.1884 (0.2640) / 0.2257 (0.3331) |
| R.m.s deviations | | | |
| Bond lengths (Å) | 0.004 | 0.003 | 0.002 |
| Bond angles (°) | 0.71 | 0.5 | 0.52 |
| Ramachandran* | | | |
| favored (%) | 98.9 | 97.2 | 97.2 |
| allowed (%) | 1.1 | 2.8 | 2.8 |
| outliers (%) | 0 | 0 | 0 |
| MolProbity Score/Percentile | 0.85 (100th) | 1.29 (100th) | 1.19 (100th) |

*calculated with MolProbity

##### Table S2. Kinase phosphorylation of EphA2 linker peptide with pS897

| Protein Kinase | CPM | Protein Kinase | CPM | Protein Kinase | CPM | Protein Kinase | CPM |
|---|---|---|---|---|---|---|---|
| PKC zeta | 192,007 | PDK1 | 10,417 | MEKK2 | 4,367 | MARK1 | 2,197 |
| PAK3 | 151,679 | CAMKK1 | 10,287 | CK2 alpha 2 | 4,254 | MNK1 | 2,187 |
| CK1 gamma 1 | 139,024 | AMPK (A1/B2/G2) | 10,227 | MLK4 | 4,176 | MEKK6 | 2,178 |
| NEK7 | 119,908 | ZAK | 10,208 | GSK3 alpha | 4,094 | STK21(CIT) | 2,124 |
| PKC beta I | 115,018 | NUAK1 | 10,059 | CLK1 | 4,068 | TAK1-TAB1 | 2,080 |
| PKC beta II | 111,980 | ALK6 (BMPR1B) | 10,006 | HIPK3 | 4,009 | ERK5 | 2,063 |
| PKC gamma | 108,805 | TSSK2 | 9,883 | VRK1 | 3,923 | DRAK2 (STK17B) | 1,980 |
| TBK1 | 106,427 | DAPK3 | 9,822 | AMPK (A2/B2/G3) | 3,912 | CLK3 | 1,958 |
| PKC alpha | 102,845 | AURORA C | 9,740 | RSK4 | 3,846 | MRCK alpha | 1,936 |
| NEK2 | 99,876 | CAMK2 gamma | 9,676 | BRAF | 3,826 | GRK6 | 1,928 |
| MLK1 | 98,444 | VRK2 | 9,550 | CDK2/CyclinA1 | 3,817 | IKK alpha | 1,852 |
| PKC eta | 95,137 | CHK1 | 9,447 | BRSK1 | 3,800 | NEK8 | 1,844 |
| PKC theta | 82,115 | STK33 | 9,190 | CASK | 3,793 | PCTK3 (CDK18) | 1,812 |
| PKC iota | 75,819 | ALK1 | 8,852 | EEF2K | 3,739 | DYRK4 | 1,805 |
| AURORA B | 72,966 | PHKG2 | 8,761 | MST4 | 3,722 | PFTK1(CDK14)Cyc | 1,776 |
| CAMK2 beta | 71,807 | CAMK1 beta | 8,751 | SNRK | 3,679 | SIK | 1,724 |
| NEK6 | 67,524 | CDK5/p25 | 8,524 | CAMK1 | 3,657 | MEKK1 | 1,691 |
| ULK1 | 65,909 | MSK1 | 8,483 | MARK3 | 3,624 | PEAK1 | 1,680 |
| ULK2 | 64,661 | PRKG1 | 8,362 | PCTK2 (CDK17) | 3,596 | PKC nu | 1,680 |
| PKC delta | 59,375 | PRKG2 | 8,340 | CDK5/p35 | 3,590 | PDHK1 | 1,668 |
| CK1 alpha 1 | 58,763 | TAOK1 | 8,257 | NLK | 3,584 | MST3 | 1,645 |
| CDC7/DBF4 | 55,210 | TGFBR2 | 8,047 | MYLK3 | 3,583 | NDR | 1,637 |
| CK1 gamma 3 | 54,848 | NIM1 | 7,945 | CAMK1 delta | 3,539 | SRPK2 | 1,632 |
| PKAc beta | 51,041 | PAK7 | 7,916 | MLCK | 3,528 | RAF1(EE) | 1,607 |
| CK1 alpha 1L | 45,567 | MEKK3 | 7,799 | RIPK1 | 3,521 | CDK2/CyclinO | 1,586 |
| CK1 gamma 2 | 40,800 | IKK beta | 7,760 | STK3 | 3,499 | p38 gamma | 1,576 |
| SGK3 | 40,181 | RSK2 | 7,721 | MYLK4 | 3,498 | ERK2 | 1,570 |
| MST1 | 38,378 | p70S6K | 7,590 | HIPK4 | 3,469 | LATS2 | 1,566 |
| TTBK2 | 35,044 | CAMKK2 | 7,398 | CLK2 | 3,416 | GRK7 | 1,543 |
| PKAc alpha | 34,771 | TLK2 | 7,236 | CAMK1 gamma | 3,353 | MEK5 | 1,504 |
| TLK1 | 34,750 | AMPK (A2/B2/G1) | 7,235 | BRSK2 | 3,331 | JNK1 | 1,491 |
| TTBK1 | 34,299 | DAPK1 | 7,166 | CDK2/CyclinE1 | 3,316 | HUNK | 1,471 |
| EIF2AK3 | 32,945 | AMPK (A1/B2/G1) | 6,987 | MAPKAPK2 | 3,270 | EIF2AK2 | 1,429 |
| PKC epsilon | 32,376 | PKD2 | 6,956 | AKT3 | 3,269 | p70S6Kb | 1,419 |
| KDR | 32,100 | AKT1 | 6,708 | MAK | 3,166 | MEK6 | 1,362 |
| GCK | 28,599 | NIK | 6,688 | PIM1 | 3,137 | PLK2 | 1,312 |
| NEK9 | 27,568 | WNK1 | 6,613 | FASTK | 3,133 | CDK6/CyclinD1 | 1,283 |
| PKAc gamma | 27,509 | MAPKAPK5 | 6,561 | BUB1B | 3,121 | GSK3 beta | 1,281 |
| PAK1 | 27,316 | MARK4 | 6,474 | COT | 3,092 | KSR1 | 1,192 |
| AMPK (A1/B1/G2) | 24,259 | CDK1/CyclinA2 | 6,408 | CDK9/CyclinK | 3,040 | MNK2 | 1,186 |
| CK1 delta | 23,576 | MYO3 alpha | 6,379 | SRPK1 | 3,030 | GRK3 | 1,154 |
| YSK4 | 23,510 | CAMK2 delta | 6,359 | ERN1 (IRE1) | 3,002 | PDHK4 | 1,153 |
| ULK3 | 21,762 | KHS1 | 6,222 | MAPKAPK3 | 2,979 | SIK3 | 1,123 |
| NEK5 | 20,936 | CDK1/CyclinA1 | 5,961 | p38 alpha | 2,945 | IKK epsilon | 1,109 |
| TAOK3 | 20,457 | ALK2 | 5,902 | TXK | 2,945 | NDR2 (STK38L) | 1,104 |
| EIF2AK4 (GCN2) | 17,969 | ALK4 | 5,897 | RIPK5 | 2,930 | DMPK | 1,103 |
| DCAMKL2 | 17,207 | AKT2 | 5,830 | ROCK1 | 2,884 | MSK2 | 1,103 |
| CK1 epsilon | 16,577 | HGK | 5,774 | PAK6 | 2,861 | Haspin (GSG2) | 1,096 |
| CDK9/CyclinT2 | 15,002 | PDHK3 | 5,753 | DYRK3 | 2,847 | PLK3 | 1,057 |
| RIPK3 | 14,926 | AMPK (A2/B1/G3) | 5,687 | HIPK2 | 2,842 | PAK2 | 1,036 |
| HPK1 | 14,096 | p38 beta | 5,478 | PIM3 | 2,757 | PDHK2 | 1,036 |
| STK32B(YANK2) | 13,961 | TAOK2 | 5,418 | SBK1 | 2,745 | ERK1 | 1,033 |
| AURORA A | 13,527 | DRAK1 (STK17A) | 5,371 | NUAK2 | 2,665 | MEK1 | 1,031 |
| AMPK (A1/B1/G3) | 13,405 | GRK1 | 5,206 | CDK7/CyclinH1/MN | 2,662 | SGK1 | 1,025 |
| DCAMKL1 | 13,380 | ALK3 (BMPR1A) | 5,101 | PIM2 | 2,623 | MEK2 | 1,009 |
| MYLK2 | 13,209 | CHK2 | 5,081 | DYRK1a | 2,620 | CDK3/CyclinE1 | 1,000 |
| PLK4 | 12,525 | AMPK (A1/B2/G3) | 5,073 | p38 delta | 2,614 | TESK2 | 973 |
| DYRK2 | 12,429 | PAK4-GTP | 5,035 | PASK | 2,590 | TGFBR1 (ALK5) | 959 |
| TTK | 12,279 | HIPK1 | 5,000 | SGK2 | 2,588 | NEK1 | 949 |
| BMPR2 | 11,689 | CLK4 | 4,997 | TSSK1B | 2,578 | LOK | 910 |
| CAMK2 alpha | 11,654 | DAPK2 | 4,974 | MLK3 | 2,548 | IRAK2 | 899 |
| NEK11 | 11,639 | TNIK | 4,961 | MELK | 2,533 | LRRK2 | 899 |
| CK2 alpha 1 | 11,509 | ERN2 (IRE2) | 4,930 | PHKG1 | 2,490 | PKN3/PRK3 | 892 |
| RSK3 | 11,244 | ASK1 | 4,850 | GRK5 | 2,471 | JNK3 | 860 |
| ROCK2 | 11,187 | IRAK4 | 4,848 | PLK1 | 2,467 | GLK | 818 |
| STK39 (STLK3) | 10,975 | PKC mu | 4,839 | GRK2 | 2,427 | PRKX | 802 |
| AMPK (A2/B1/G2) | 10,972 | CDK6/CyclinD3 | 4,781 | RIPK2 | 2,371 | LATS1 | 797 |
| AMPK (A2/B1/G1) | 10,963 | SLK | 4,666 | QIK | 2,354 | STK36 | 794 |
| PKN2/PRK2 | 10,961 | TOPK | 4,627 | EIF2AK1 (HRI) | 2,353 | STK19 | 793 |
| RSK1 | 10,918 | CAMK4 | 4,553 | MINK1 | 2,318 | JNK2 | 751 |
| NEK3 | 10,845 | MYO3 beta | 4,527 | MSSK1 | 2,248 | CDK4/CyclinD1 | 745 |
| AMPK (A2/B2/G2) | 10,827 | CDK2/CyclinA2 | 4,455 | ICK | 2,241 | STK32C (YANK3) | 685 |
| AMPK (A1/B1/G1) | 10,681 | MARK2 | 4,441 | KSR2 | 2,236 | PKN1/PRK1 | 51 |
| MLK2 | 10,481 | NEK4 | 4,413 | PCTK1(CDK16)Cyc | 2,226 | | |
| SOK1 | 10,460 | MRCK beta | 4,372 | LIMK1 | 2,221 | | |

| |
|---|
| > 100,000 |
| 50,000 - 99,999 |
| 25,000 - 49,999 |
| 10,000 - 24,999 |

#### SUPPLEMENTARY FIGURE LEGENDS

**Figure S1. EphA2 is phosphorylated on multiple serine/threonine residues in the kinase-SAM linker in cancer cell lines.** (**A**) Number of phosphopeptides identified by mass spectrometry as containing one of the five phosphosites indicated. The somewhat higher abundance of peptides phosphorylated on S897 may be due to the many experiments in which peptides were purified for analysis using phospho-AKT substrate antibodies, which recognize the phosphorylated S897 motif. Data are from PhosphoSite (phosphosite.org). (**B, C**) A number of cancer cell lines were screened in two independent experiments for EphA2 phosphorylation on S892, S897 and S901. EphA2 immunoprecipitates were probed with an antibody specific for pS892 and reprobed for EphA2. Lysates were probed in separate blots with antibodies specific for pS897 or pS901 and reprobed for EphA2. Amido black staining of the region of the blot between ~65 kDa and ~120 kDa shows relative protein loading in different lanes. The white vertical line in C indicates removal of an irrelevant lane. (**D**) Correlation of pS892 or pS901 versus pS897, normalized to total EphA2 levels, in the experiments shown in C and D. The values in each experiment were normalized to the value for PC3 cells (in red) measured in the same experiment. (**E**) The number of peptides containing each phosphosite, identified by mass spectrometry in lung and breast cancer cell lines, were normalized to the number of peptides containing the same phosphorylation site in a reference pool of 16 non-small cell lung cancer cell lines. The original data used for the analyses show are from ref.[30]. In E and F, the Pearson correlation coefficient, r, and the P value for the significance of the correlation are shown. The best fit interpolation line (thick black line) and 95% confidence interval (thin grey dotted lines) are shown in the panels with significant correlation. (**F**) The majority (62%) of the phosphorylated peptides from the EphA2 linker, identified by mass spectrometry, contain multiple (two or three) phosphorylation sites. Data are from the PhosphoSite database (phosphosite.org).

**Figure S2. The EphA2 intracellular region is monomeric in solution.** Sedimentation velocity measured by analytical ultracentrifugation for the EphA2 WT intracellular region, showing that it is a monomer with an apparent molecular weight of 42.6 kDa, which closely matches the calculated molecular weight of 43.9 kDa.

**Figure S3. Molecular details of the interaction between the non-phosphorylated EphA2 juxtamembrane segment and the kinase domain.** (**A**) In our structure (PDB: 7KJA), the EphA2 juxtamembrane segment (molecule A shown) forms a short, 2-turn helix comprising residues P597–F604. Y594 is surrounded by a hydrophobic pocket lined by I666 and F670 at the end of the αC-helix and residues M733 and Y735, and additionally forms a hydrogen bond with the backbone of M733 (dashed line). (**B**) In a previous structure of the EphA2 juxtamembrane segment and kinase domain (residues K586-I875; PDB 5EK7[37]), Y588 rather than Y594 binds to the same hydrophobic pocket. In this previous structure, the juxtamembrane segment forms an additional short helix (outlined by a dotted green line) leading to the observed differences. (**C**) Superposition of molecules A and B of our EphA2 WT structure (colors as in Fig. 1), an EphA2 kinase domain structure (residues A599-P896; PDB 6Q7D, unpublished, grey) and an EphA3 kinase domain (PDB 2QOC[38], orange). Linker residues I875-T898 of the two molecules in our EphA2 structure have the same conformation, which differs from the linker conformations of EphA2 without the SAM domain, whereas some features (N-terminal part of the linker and interactions of the I893 hydrophobic residue) resemble the linker conformation of EphA3 without the SAM domain. Kinase domain residues forming a hydrophobic pocket that accommodates EphA2 I893 or EphA3 L901 are shown as sticks. (**D**) The two molecules in the EphA2 WT structure were superposed based on their SAM domains. The first residue of the SAM domain (F906) has the same position in the two EphA2 molecules, whereas differences are observed for the first residue after the missing portion of the linker (P905). Colors as in Fig. 1.

**Figure S4. Crystal structure of the intracellular region of the EphA2 S901E mutant.** Superposition of the structures of the intracellular regions of EphA2 WT (colored as in Fig. 1) and the S901E mutant (grey). The two structures are almost identical, with an overall RMSD calculated by aligning all Cα atoms of 0.48 Å. The RMSD obtained by aligning the individual molecules is 0.37 Å for molecule A and 0.34 Å for molecule B. A notable difference between the two structures is that most of the activation loop (residues L760-I779) is undefined in the EphA2 WT structure (the undefined region of the loop is not shown), whereas only a small portion of the activation loop (T773-G777) is undefined in molecule A of the EphA2 S901E mutant structure (indicated by a dashed line). This difference is possibly due to the slightly smaller unit cell dimensions and tighter crystal packing of the S901E mutant. The inset shows an enlargement of the indicated portion of the structures in a slightly tilted orientation. AMPPcP is shown as sticks, $Mg^{2+}$-ions as purple spheres.

**Figure S5. Time course of hydrogen-deuterium exchange for key peptides.** Deuterium uptake in the indicated peptides is plotted against time. The Y-axes are scaled to the maximum theoretically possible deuterium uptake for each peptide. Colors denote different mutants, as indicated. Averages +/- SEM from 3 technical replicates are shown.

**Figure S6. Coverage plots from HDX-MS experiments and relative fractional uptake after 0.5 min deuterium exposure.** (**A**) EphA2 WT (**B**) 5E mutant (**C**) 3E mutant (**D**) S892E mutant (**E**) S897E mutant (**F**) E820K/E825K mutant. In six experiments, we assigned 111 peptides covering 97.8% of the EphA2 WT sequence with 4.0 average redundancy.

**Figure S7. Putative EphA2 dimer.** (**A**) Crystallographic EphA2 kinase domain dimer (PDB: 6FNG)[93] with electrostatic surface. (**B**) Open book view of the dimer shown in A. The dimer interfaces are outlined by green dotted lines. (**C**) View as in B, both molecules are colored in wheat with dimer interfaces shown in cyan.

**Figure S8. FSI-FRET analysis of dimerization for EphA2 kinase-SAM linker mutants.** (**A**) FRET efficiencies, measured in individual cells, are shown as a function of EphA2-EYFP (acceptor) concentrations in the first and third columns. The data for the EphA2 phosphomimetic mutants (red symbols) and non-phosphorylatable mutants (blue symbols) are compared to EphA2 WT data (black symbols; from ref.[94]). These FRET efficiencies have been corrected as described[95] for "proximity FRET", which arises due to random proximity of donors and acceptors in the two-dimensional plasma membrane. Best fit dimerization curves were obtained by fitting the FRET data with equation (3) in the Methods and are shown as solid lines in the second and fourth columns. The experimental data were binned and are shown as averages and standard errors. The number of data points, N, is as follows: WT (N=670), 5E (N=314), 3E (N=235), S897E (N=191), S892E (N=188), 5A (N=234), ASAAA (N=317), S897A (N=196), and S892A (N=241). (**B**) Best-fit parameters. $K_{diss}$ is the two-dimensional dissociation constant. The errors are 68% confidence intervals from the fit. Statistical significance ($P<0.05$) for comparison with EphA2 WT was determined by one-way ANOVA followed by the Tukey's multiple comparisons test and is indicated with an asterisk. $\tilde{E}$ is the Intrinsic FRET, which depends on the positioning of the fluorescent proteins in the EphA2 dimers. The average distance between the fluorescent proteins in the EphA2 dimers, $d$, is calculated from $\tilde{E}$ using equation (2), under the assumption of free rotation of the fluorescent proteins.

**Figure S9. Hypothetical model illustrating the proposed effects of phosphorylation on the arrangement of the EphA2 intracellular region.** (**A**) In the unphosphorylated, inactive EphA2 intracellular region, an electronegative surface of the kinase domain interacts with an electropositive surface of the SAM domain (SAM). In addition, the C-terminal portion of the juxtamembrane segment (JXTM) interacts with the kinase domain and its N-terminal portion together with an electropositive surface of the kinase domain mediate association with the plasma membrane. (**B**) Phosphorylation of serine/threonine residues in the kinase-SAM linker promotes transition to an open conformation. Surfaces of the kinase and/or SAM domain exposed in the open conformation may become available to interact with potential binding partners mediating non-canonical signaling responses. (**C**) Tyrosine phosphorylation of two conserved tyrosines in the EphA2 juxtamembrane segment and other tyrosines disrupts the interaction with the kinase domain and promote distancing of the kinase domain from the plasma membrane. Tyrosine phosphorylated motifs in the juxtamembrane segment also bind SH2 domain-containing effectors that mediate canonical signaling. The black double-headed arrow indicates a potential interplay of canonical and non-canonical signaling mediated by an intramolecular allosteric network coupling the EphA2 linker, juxtamembrane segment and kinase domain. In addition, EphA2 canonical signaling can inhibit non-canonical signaling in some cellular contexts through inhibition of AKT or ERK, as indicated by the blue inhibitory arrow.

**Figure S10. EphA2 W819 and E820 as sensors of kinase-SAM linker phosphorylation.** Overlay of the EphA2 WT kinase-SAM domain structure (molecule B), the EphA2 S897E/S901E mutant kinase-SAM domain structure, and the EphA3 kinase-linker structure (PDB 2QOC[38]). In EphA2, E820 in the αFG-loop is positioned just above S892 and may act as a sensor for linker phosphorylation. The EphA2 αFG-loop, including W819, is part of an allosteric network that relays information about linker phosphorylation to the activation loop. The E827 sidechain is observed in two different conformations in the EphA3 crystal structure.

### Figure S1

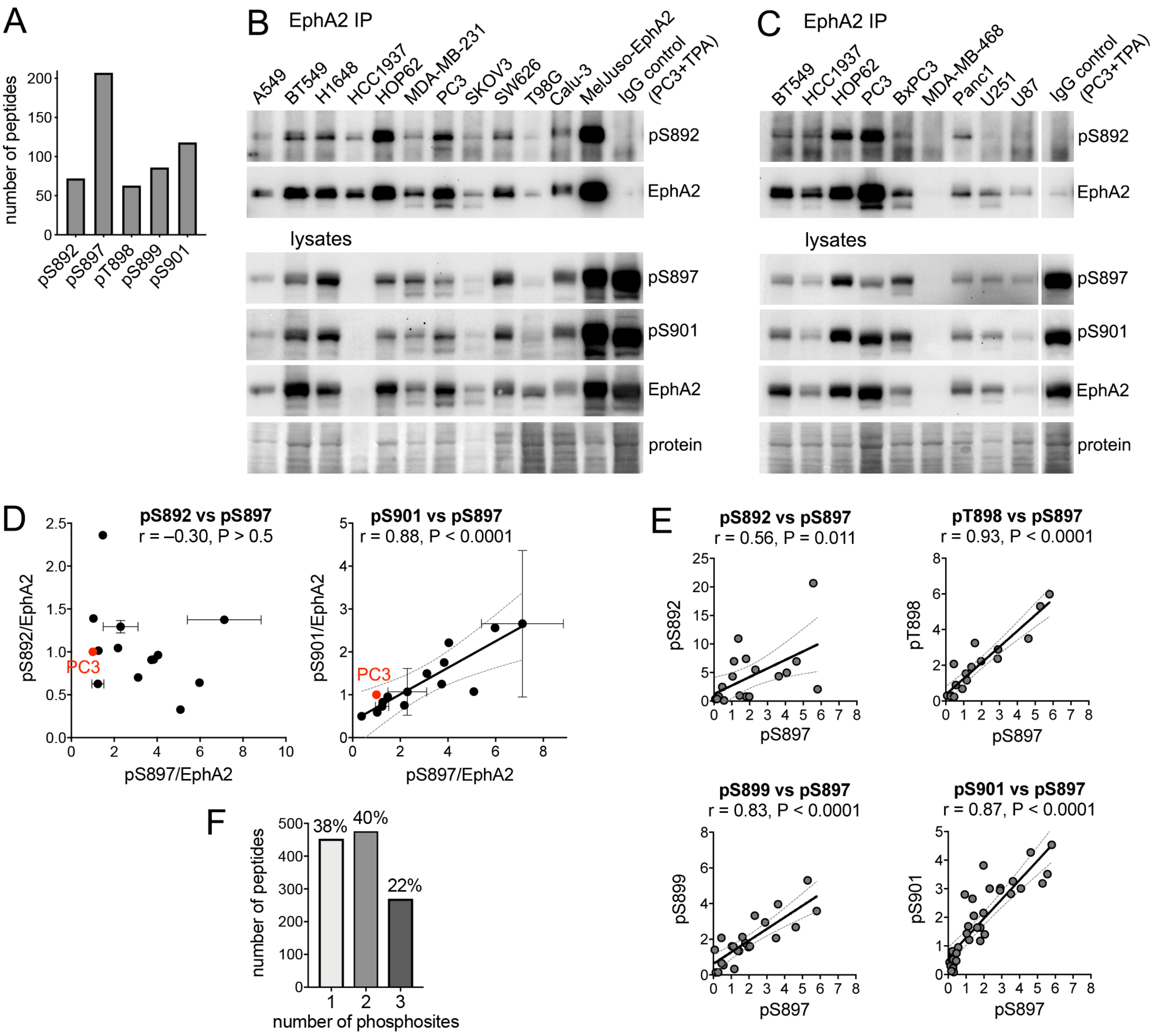

A
number of peptides
pS892
pS897
pT898
pS899
pS901
B
EphA2 IP
A549
BT549
H1648
HCC1937
HOP62
MDA-MB-231
PC3
SKOV3
SW626
T98G
Calu-3
MelJuso-EphA2
IgG control (PC3+TPA)
pS892
EphA2
lysates
pS897
pS901
EphA2
protein
C
EphA2 IP
BT549
HCC1937
HOP62
PC3
BxPC3
MDA-MB-468
Panc1
U251
U87
IgG control (PC3+TPA)
pS892
EphA2
lysates
pS897
pS901
EphA2
protein
D
pS892 vs pS897
r = –0.30, P > 0.5
pS892/EphA2
pS897/EphA2
PC3
pS901 vs pS897
r = 0.88, P < 0.0001
pS901/EphA2
pS897/EphA2
PC3
E
pS892 vs pS897
r = 0.56, P = 0.011
pS892
pS897
pT898 vs pS897
r = 0.93, P < 0.0001
pT898
pS897
pS899 vs pS897
r = 0.83, P < 0.0001
pS899
pS897
pS901 vs pS897
r = 0.87, P < 0.0001
pS901
pS897
F
number of peptides
38%
40%
22%
number of phosphosites

Figure S2

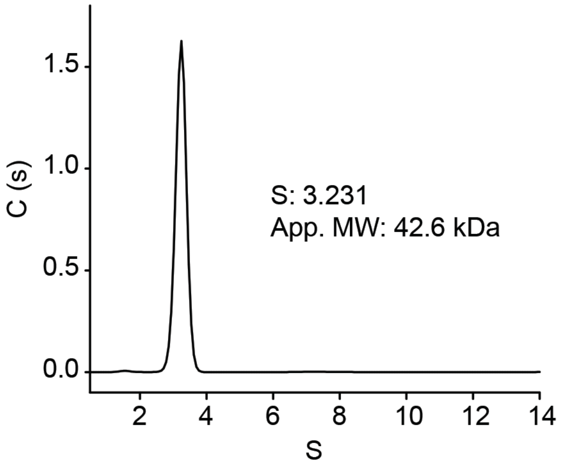

### Figure S3

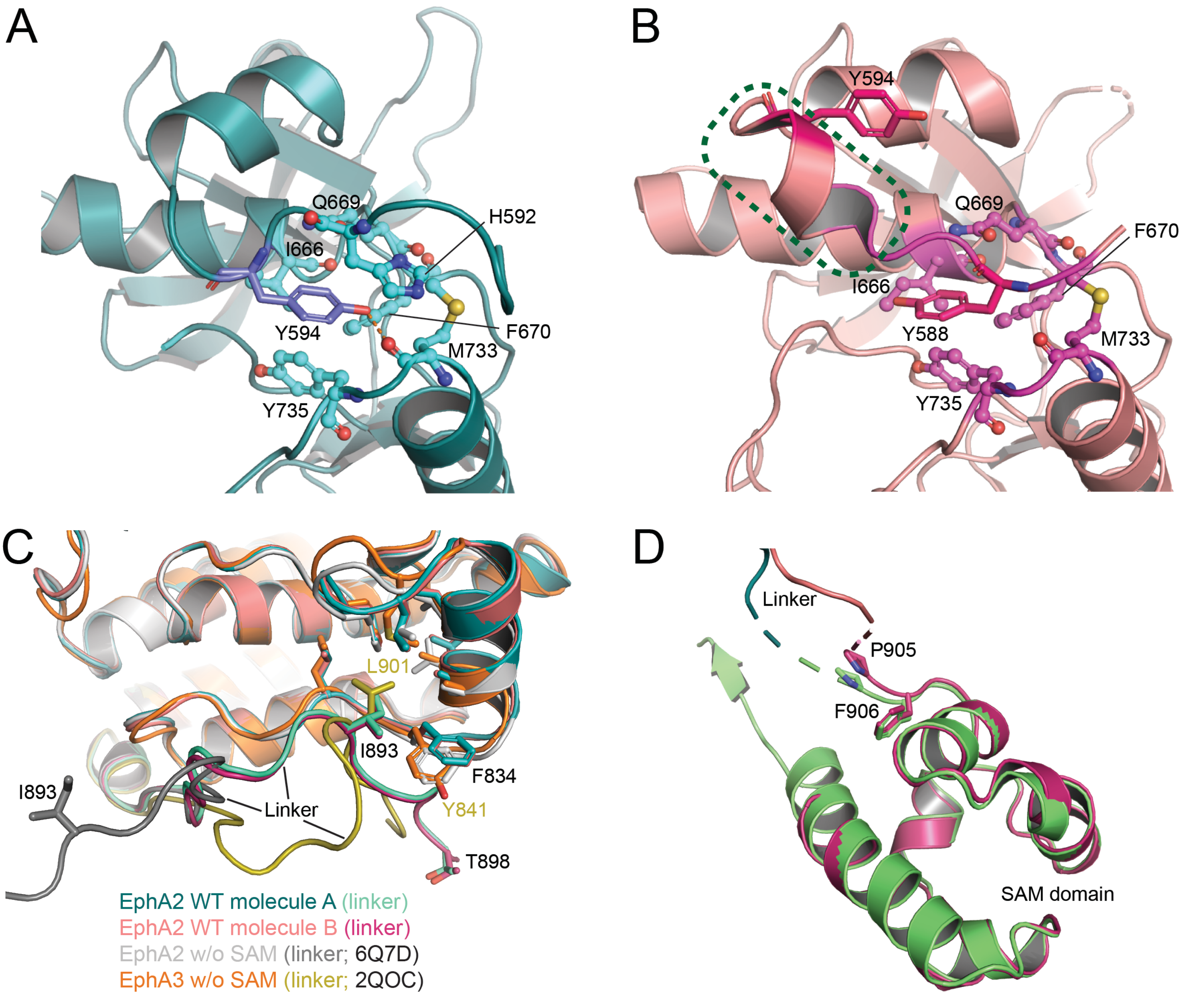
A
Q669
H592
I666
Y594
F670
M733
Y735
B
Y594
Q669
F670
I666
Y588
M733
Y735
C
L901
I893
F834
I893
Linker
Y841
T898
EphA2 WT molecule A (linker)
EphA2 WT molecule B (linker)
EphA2 w/o SAM (linker; 6Q7D)
EphA3 w/o SAM (linker; 2QOC)
D
Linker
P905
F906
SAM domain

### Figure S4

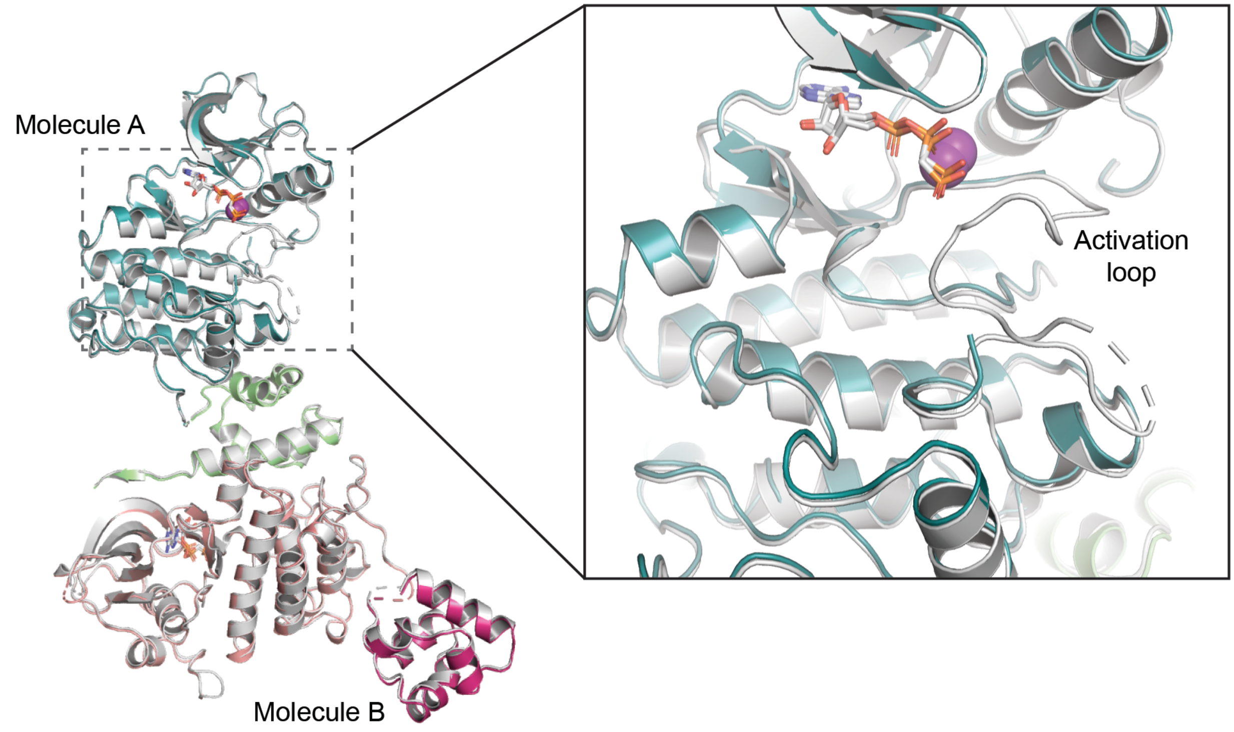

Figure S5

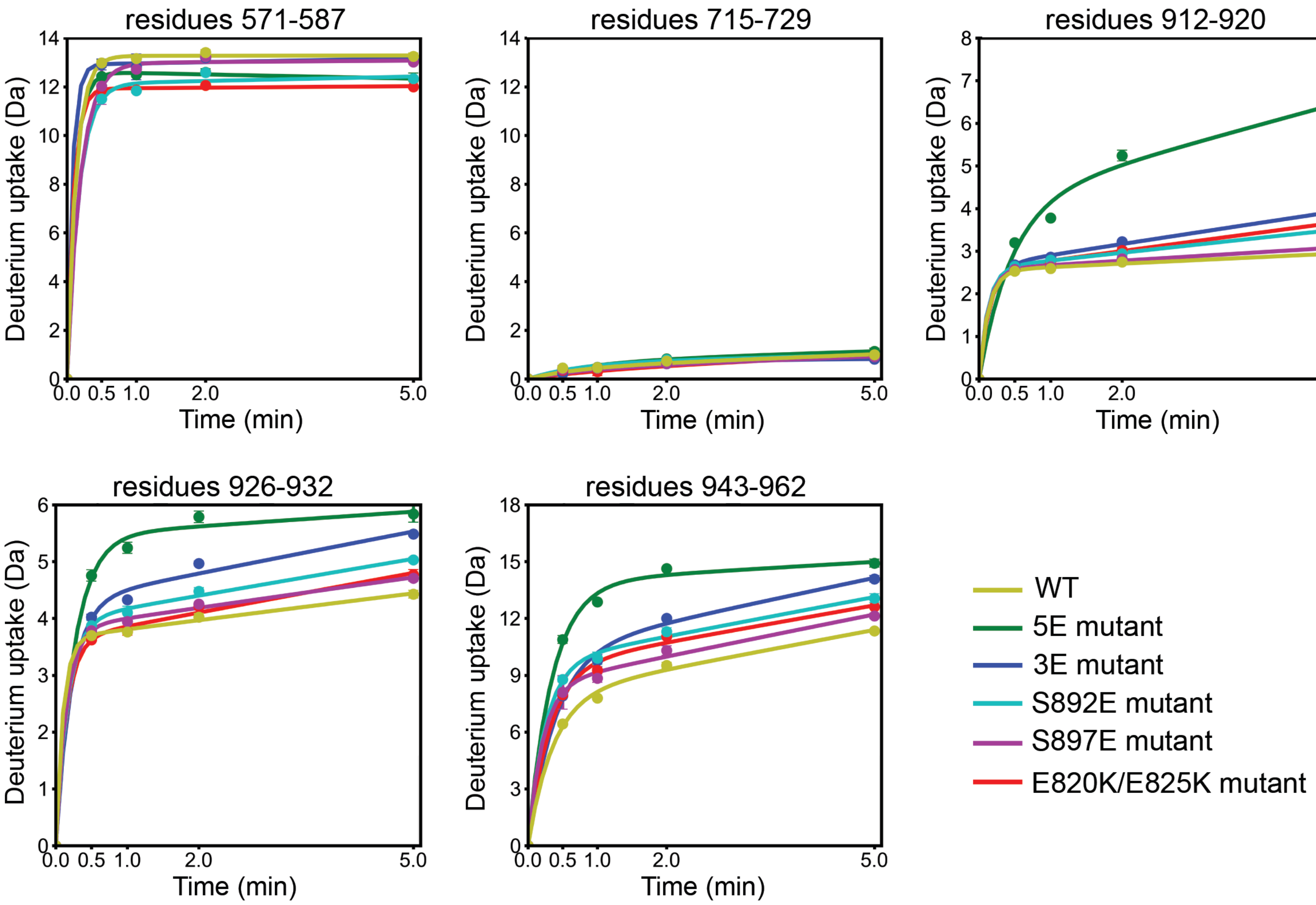

residues 571-587
residues 715-729
residues 912-920
residues 926-932
residues 943-962
Deuterium uptake (Da)
Time (min)
0.0 0.5 1.0 2.0 5.0
WT
5E mutant
3E mutant
S892E mutant
S897E mutant
E820K/E825K mutant

Figure S6

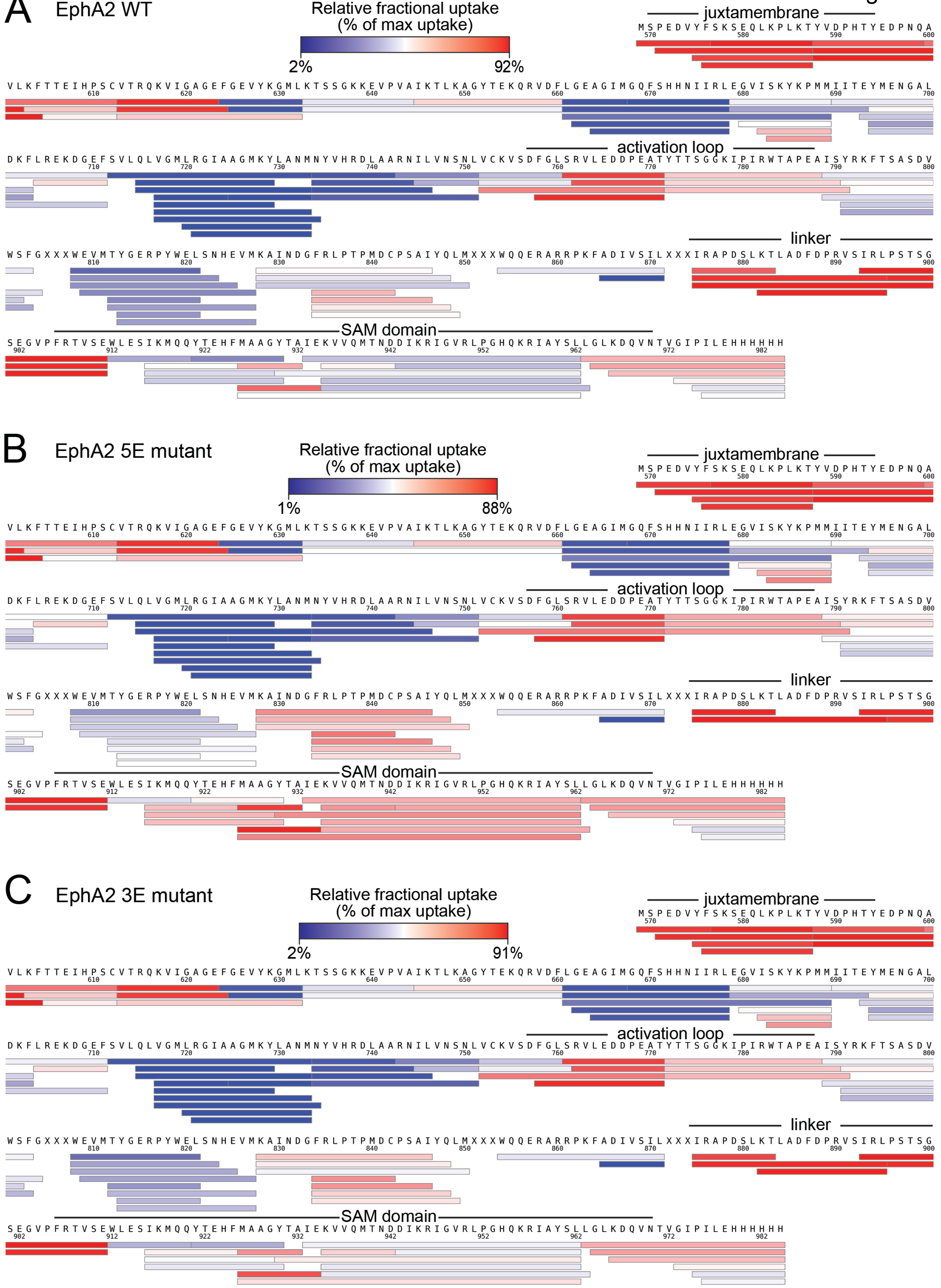

A
EphA2 WT
Relative fractional uptake (% of max uptake)
2%
92%
juxtamembrane
activation loop
linker
SAM domain
B
EphA2 5E mutant
Relative fractional uptake (% of max uptake)
1%
88%
juxtamembrane
activation loop
linker
SAM domain
C
EphA2 3E mutant
Relative fractional uptake (% of max uptake)
2%
91%
juxtamembrane
activation loop
linker
SAM domain

Figure S6

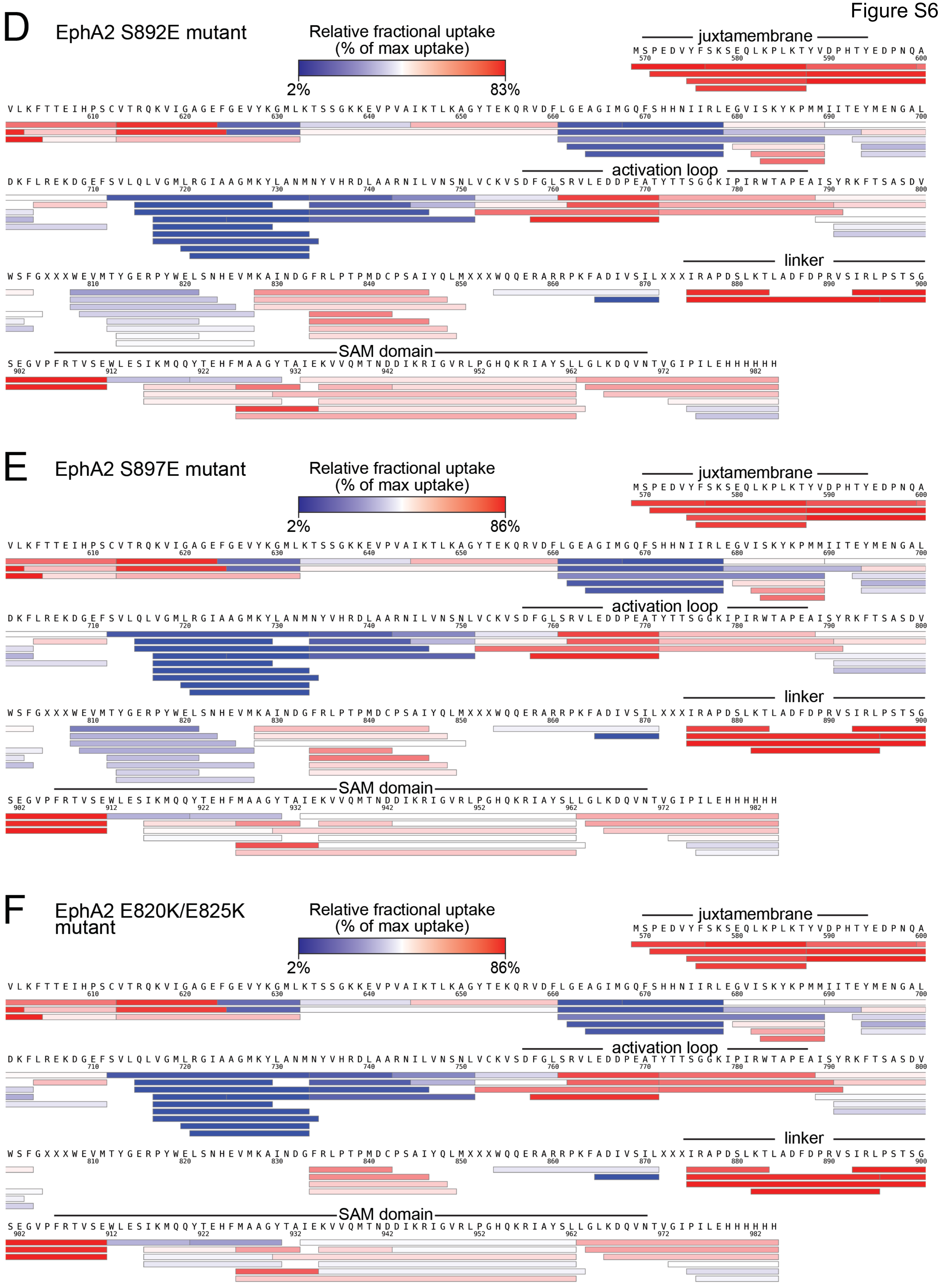
D
EphA2 S892E mutant
Relative fractional uptake
(% of max uptake)
2%
83%
juxtamembrane
activation loop
linker
SAM domain
E
EphA2 S897E mutant
Relative fractional uptake
(% of max uptake)
2%
86%
juxtamembrane
activation loop
linker
SAM domain
F
EphA2 E820K/E825K mutant
Relative fractional uptake
(% of max uptake)
2%
86%
juxtamembrane
activation loop
linker
SAM domain

Figure S7

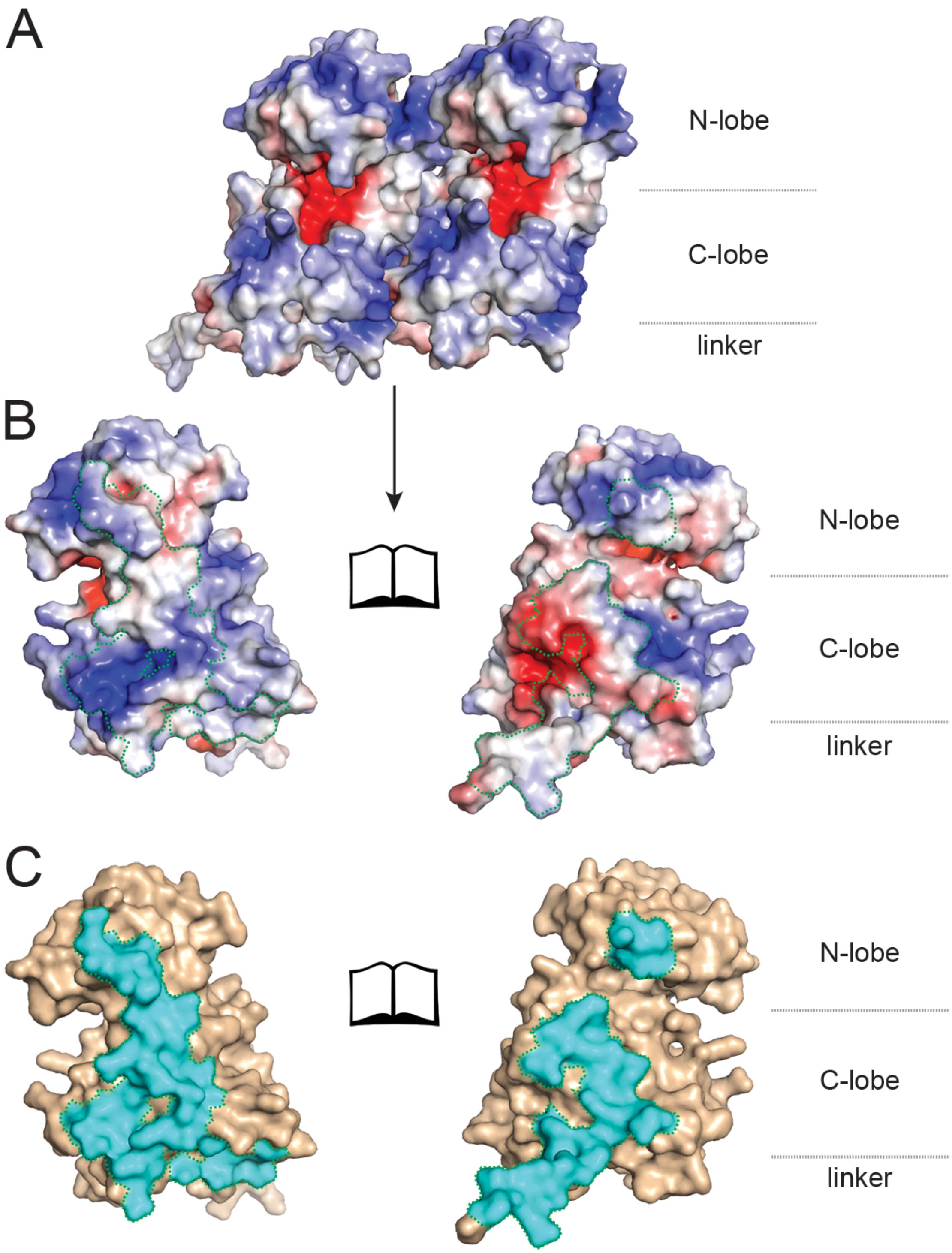

A
N-lobe
C-lobe
linker
B
N-lobe
C-lobe
linker
C
N-lobe
C-lobe
linker

### Figure S8

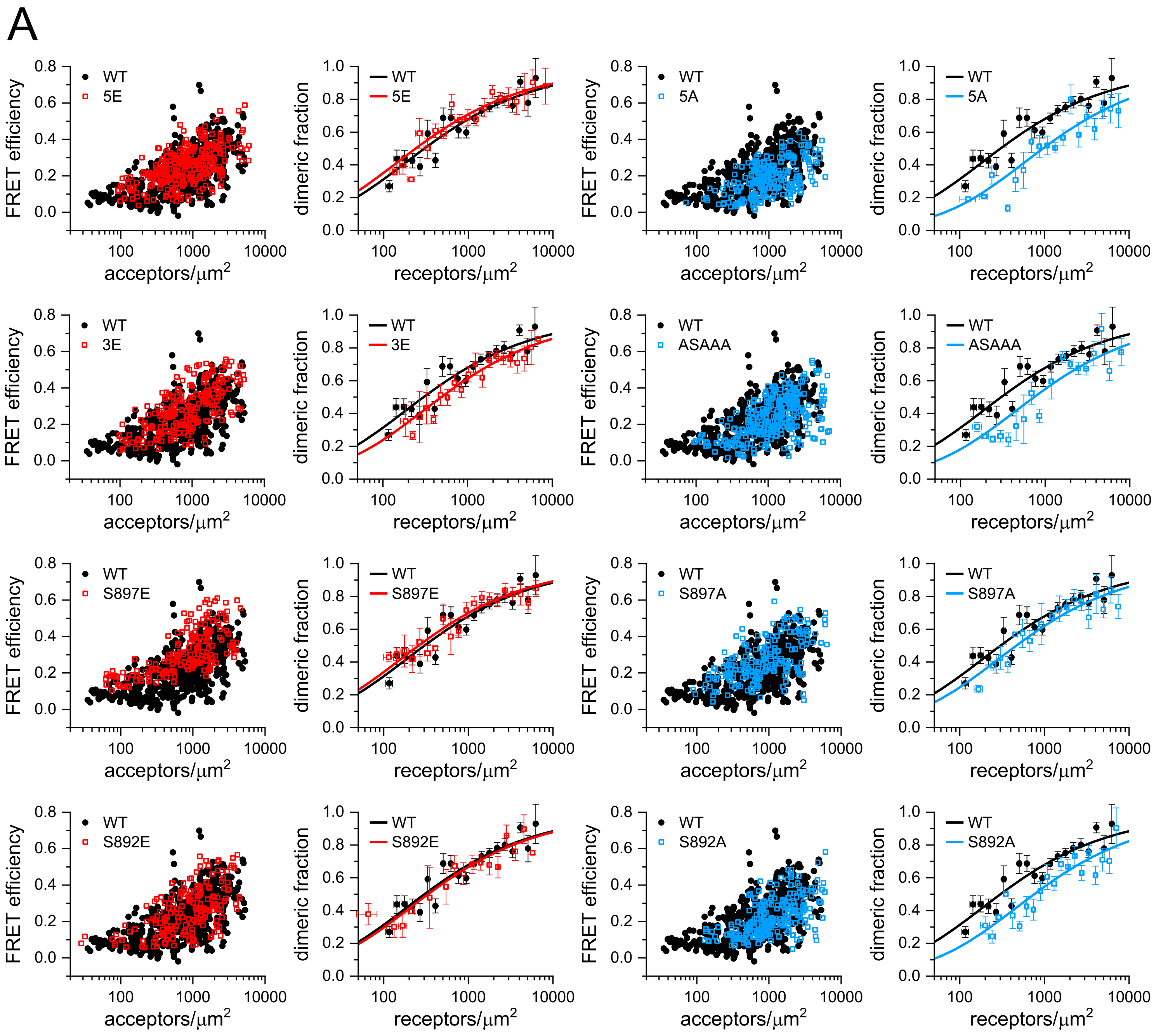


B

| EphA2 | $K_{diss}$ (rec/μm$^2$) | Ẽ | d (Å) |
|---|---|---|---|
| WT | 302 ± 68 | 0.53 ± 0.02 | 53.6 ± 0.7 |
| 5E | 238 ± 73* | 0.53 ± 0.03 | 53.4 ± 0.9 |
| 3E | 409 ± 144* | 0.69 ± 0.03 | 47.6 ± 1.3 |
| S897E | 259 ± 63 | 0.76 ± 0.03 | 44.8 ± 1.2 |
| S892E | 334 ± 154 | 0.58 ± 0.04 | 51.7 ± 1.6 |
| 5A | 958 ± 347* | 0.50 ± 0.04 | 54.6 ± 1.6 |
| ASAAA | 698 ± 209* | 0.55 ± 0.03 | 52.7 ± 1.3 |
| S897A | 465 ± 149* | 0.70 ± 0.04 | 47.3 ± 1.7 |
| S892A | 756 ± 247* | 0.55 ± 0.04 | 52.8 ± 1.4 |

### Figure S9

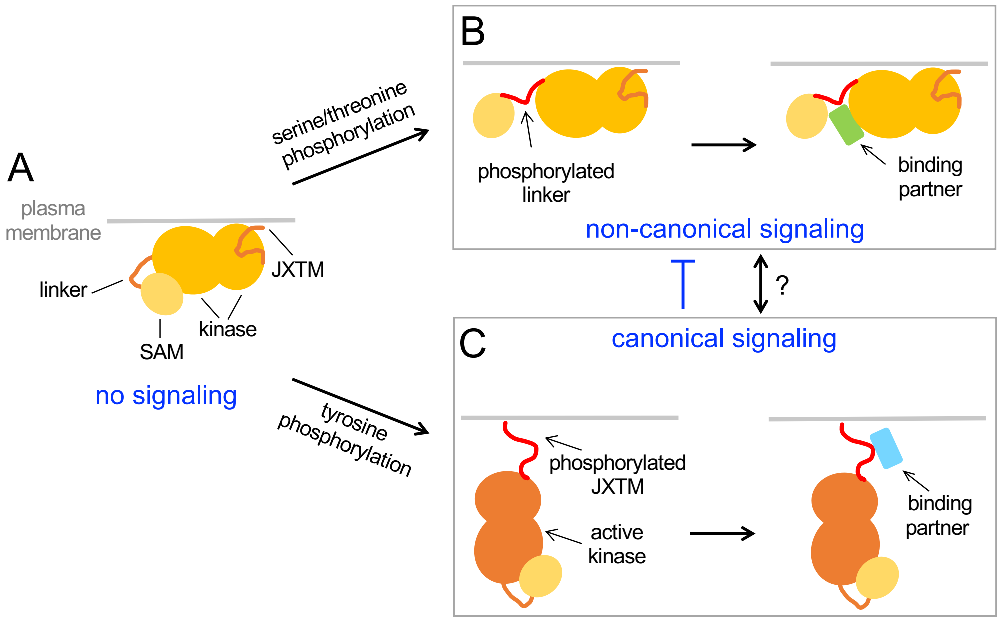
A
plasma
membrane
linker
JXTM
kinase
SAM
no signaling
serine/threonine
phosphorylation
tyrosine
phosphorylation
B
phosphorylated
linker
binding
partner
non-canonical signaling
?
C
canonical signaling
phosphorylated
JXTM
active
kinase
binding
partner

### Figure S10

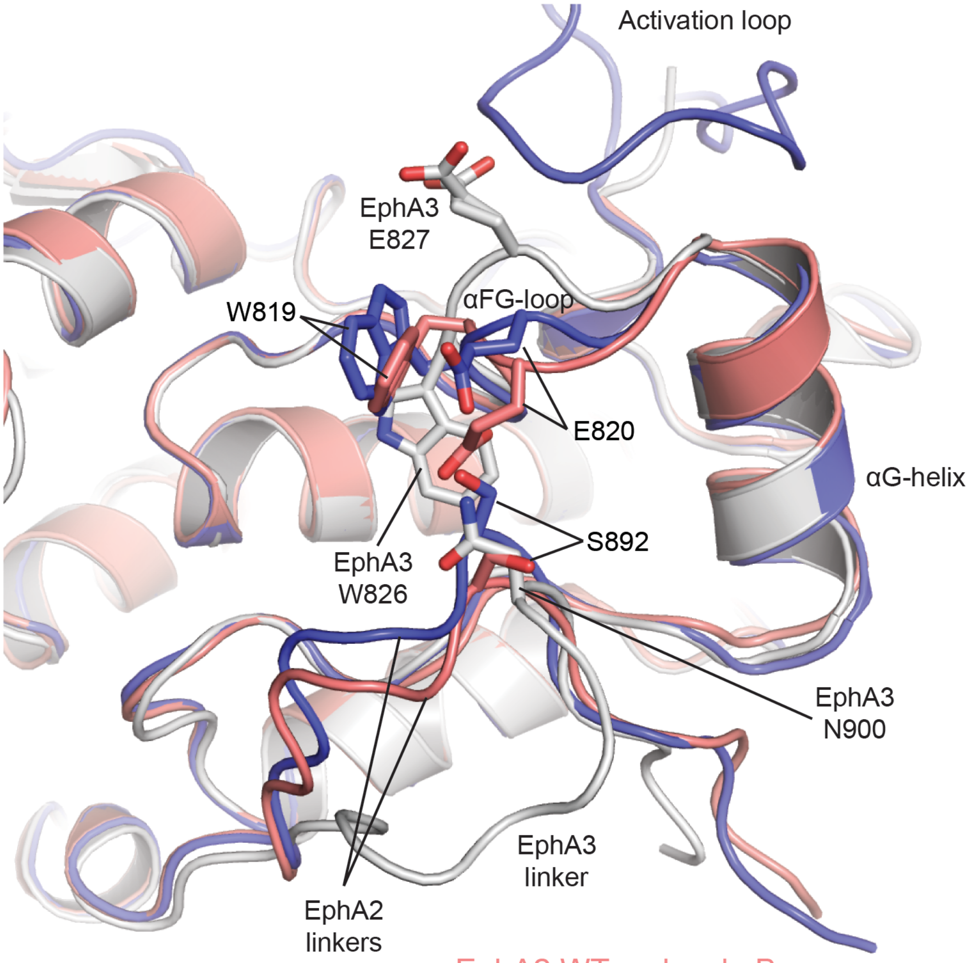
Activation loop
EphA3
E827
W819
αFG-loop
E820
αG-helix
S892
EphA3
W826
EphA3
N900
EphA3
linker
EphA2
linkers
EphA2 WT molecule B
EphA2 S897E/S901E
EphA3 w/o SAM (2QOC)
